# Induction and inhibition of *Drosophila* X chromosome gene expression are both impeded by the dosage compensation complex

**DOI:** 10.1101/2021.11.22.469548

**Authors:** Richard P. Meisel, Danial Asgari, Florencia Schlamp, Robert L. Unckless

**Author notes:** To whom correspondence should be addressed: Richard P. Meisel 3455 Cullen Blvd. Houston, TX 77204 USA.

## Abstract

Sex chromosomes frequently differ from the autosomes in the frequencies of genes with sexually dimorphic or tissue-specific expression. Multiple hypotheses have been put forth to explain the unique gene content of the X chromosome, including selection against male-beneficial X-linked alleles, expression limits imposed by the haploid dosage of the X in males, and interference by the dosage compensation complex (DCC) on expression in males. Here, we investigate these hypotheses by examining differential gene expression in *Drosophila melanogaster* following several treatments that have widespread transcriptomic effects: bacterial infection, viral infection, and abiotic stress. We found that genes that are induced (up-regulated) by these biotic and abiotic treatments are frequently under-represented on the X chromosome, but so are those that are repressed (down-regulated) following treatment. We further show that whether a gene is bound by the DCC in males can largely explain the paucity of both up- and down-regulated genes on the X chromosome. Specifically, genes that are bound by the DCC, or close to a DCC high-affinity site, are unlikely to be up- or down-regulated after treatment. This relationship, however, could partially be explained by a correlation between differential expression and breadth of expression across tissues. Nonetheless, our results suggest that DCC binding, or the associated chromatin modifications, inhibit both up- and down-regulation of X chromosome gene expression within specific contexts. We propose multiple possible mechanisms of action for the effect, including a role of Males absent on the first (Mof), a component of the DCC, as a dampener of gene expression variance in both males and females. This effect could explain why the *Drosophila* X chromosome is depauperate in genes with tissue-specific or induced expression, while the mammalian X has an excess of genes with tissue-specific expression.

## INTRODUCTION

Many animal species, as well as some plants and other eukaryotes, have sex chromosomes, which are often under different transcriptional regulation than the autosomes. Sex chromosomes can be grouped into several different categories, with XY and ZW systems amongst the most common in animals (Bachtrog *et al*. 2014). X and Z chromosome gene expression can be controlled by transcriptional regulators and histone modifications that are unique from the autosomes (Lucchesi *et al*. 2005; Ferrari *et al*. 2014; Gu *et al*. 2019). For example, one copy of the mammalian X chromosome is silenced (via the recruitment of facultative heterochromatin) in somatic tissues of XX females by a combination of non-coding RNAs and proteins (Lyon 1961; Brown *et al*. 1991; Chow *et al*. 2005). In contrast, the *Drosophila* dosage compensation complex (DCC) up-regulates gene expression on the X chromosome in males using a combination of RNAs and proteins (Lucchesi and Kuroda 2015). The DCC only assembles in male somatic tissues, where it initiates the acetylation of lysine 16 in histone H4 (H4K16ac) specifically on the X chromosome, compensating for the haploid dose (Gelbart *et al*. 2009). Furthermore, in the male germline of some animal species, there is evidence for silencing of the single X chromosome (Lifschytz and Lindsley 1972; Turner 2007), although the extent of this meiotic sex chromosome inactivation (MSCI) varies across taxa (Bean *et al*. 2004; Meiklejohn *et al*. 2011; Turner 2015).

The unique transcriptional and chromatin environments of X chromosomes, along with their hemizygosity in males, create selection pressures on X-linked genes that differ from the autosomes, resulting in X-autosome differences in gene content that are taxon-specific. For example, the mammalian X chromosome is enriched for genes that are expressed specifically in male reproductive tissues, such as the prostate and testis (Wang *et al*. 2001; Lercher *et al*. 2003; Mueller *et al*. 2008, 2013; Meisel *et al*. 2012a). In contrast, the *Drosophila melanogaster* X chromosome contains very few genes that are expressed primarily in the male-specific accessory gland, a reproductive organ analogous to the mammalian prostate (Swanson *et al*. 2001; Ravi Ram and Wolfner 2007; Meisel *et al*. 2012a). The *Drosophila* X chromosome also contains a paucity of genes with male-biased expression (i.e., up-regulated in males relative to females) relative to the autosomes (Parisi *et al*. 2003; Sturgill *et al*. 2007). Taxon-specific X-autosome differences in gene content further extend to genes with non-reproductive functions. In *D. melanogaster*, for instance, the X chromosome is deficient for genes that have narrow expression in non-reproductive tissues, whereas the mammalian X is enriched for genes with tissue-specific expression (Lercher *et al*. 2003; Mikhaylova and Nurminsky 2011; Meisel *et al*. 2012a).

Multiple hypotheses have been proposed to explain the differences in gene content between X chromosomes and autosomes (Table 1). One of these hypotheses is based upon the prediction that sexually antagonistic selection will favor recessive male-beneficial mutations (or dominant female-beneficial alleles) on the X chromosome (Rice 1984; Charlesworth *et al*. 1987). This sexual antagonism hypothesis has numerous limitations (Fry 2010), including the inability to explain differences between *Drosophila* and mammalian X chromosomes in their deficiency or enrichment, respectively, of genes expressed in male reproductive tissues (Meisel *et al*. 2012a). A second hypothesis focuses specifically on the male germline, where MSCI silences the X chromosome (Lifschytz and Lindsley 1972) and may favor duplication of genes to the autosomes (Betrán *et al*. 2002; Emerson *et al*. 2004; Potrzebowski *et al*. 2008; Vibranovski *et al*. 2009). However, there is not a deficiency of testis-biased genes on the *D. melanogaster* X chromosome (Meisel *et al*. 2012a; Meiklejohn and Presgraves 2012), limiting the ability of MSCI to explain the unique gene content of the *Drosophila* X chromosome. Third, the haploid dose of the X in males may impose a maximal gene expression level lower than the autosomes, selecting against X-linked genes with high expression (Wolfner *et al*. 1997; Vicoso and Charlesworth 2009; Hurst *et al*. 2015). This “dosage limit” hypothesis may even apply in species where the haploid X is dosage compensated by up-regulation of X-linked expression. For example, in *D. melanogaster* there may be a transcriptional limit beyond which expression cannot be exceeded or some genes may not be dosage compensated in males (Meisel *et al*.

**Table 1.** Hypotheses to explain the unique gene content of the Drosophila X chromosome.

| Hypothesis | Predicted X-autosome differences |
| --- | --- |
| Sexual antagonism | Excess or deficiency of X-linked genes with sex-specific functions and/or expressed in reproductive tissues |
| MSCI | Deficiency of X-linked testis-expressed genes |
| Dosage limit | Deficiency of highly expressed or induced X-linked genes |
| DCC-interference | Deficiency of DCC-bound genes highly expressed in males |
| Variance dampening | Deficiency of X-linked genes up- or down-regulated in specific contexts (e.g., tissues, infections, abiotic treatments) |

2012a).

Here, we focus on the effect of the DCC on X chromosome expression and gene content in *D. melanogaster*. The DCC most strongly binds to more than 100 so-called chromatin entry or high affinity sites (HAS), from which it is thought to spread across the X chromosome (Kelley *et al*. 1999; Alekseyenko *et al*. 2008; Straub *et al*. 2008). Bachtrog *et al*. (2010) observed that genes near an HAS or bound by the DCC are less likely to have male-biased expression, and genes further from an HAS have a larger magnitude of male-biased expression. This led them to hypothesize that the DCC interferes with acquisition of male-biased expression on the X chromosome.

There is mixed evidence for the hypothesis that DCC-interference is responsible for the unique gene content of the X chromosome. Consistent with the DCC-interference hypothesis, when Belyi *et al*. (2020) measured expression of a reporter construct that was inserted at random locations on the X chromosome, they found reduced expression in male somatic tissues for transgenes inserted at chromosomal loci closer to endogenous DCC binding sites. However, when genes with testis-biased expression are excluded or when somatic tissues are analyzed separately, there is no relationship between male-biased expression and distance from an HAS for endogenous genes (Vensko and Stone 2014; Gallach and Betrán 2016). In addition, genes with male-biased expression in brain or head are over-represented on the *D. melanogaster* X chromosome and closer to DCC binding sites (Huylmans and Parsch 2015), which is opposite of what is predicted by the DCC-interference hypothesis.

Our analysis addresses a fifth hypothesis, specifically whether the *Drosophila* DCC creates an unfavorable environment for X-linked genes that are differentially expressed in specific contexts. The *D. melanogaster* X chromosome is depauperate in genes with narrow expression in specific tissues (Mikhaylova and Nurminsky 2011; Meisel *et al*. 2012a), and X-linked genes further from an HAS or not bound by the DCC have more tissue-specific expression (Meisel *et al*. 2012b). In contrast, X-linked genes with female-biased expression, which also tend to be broadly expressed (Meisel 2011), are more likely to be bound by the DCC (Gallach and Betrán 2016). This suggests that the DCC creates an unfavorable environment for X-linked genes that are up- or down-regulated in specific tissues, possibly because the DCC prevents the differential regulation of gene expression across contexts. Consistent with this hypothesis, there is evidence that Mof, one of the proteins in the DCC, dampens transcriptional variation on the *D. melanogaster* X chromosome (Lee *et al*. 2018). Moreover, genes that are bound by the DCC have less genetic variation for gene expression than X-linked unbound genes (Meisel *et al*. 2012b), and transgenes inserted on the *D. melanogaster* X chromosome have less intra-locus expression variation in males than females (Belyi *et al*. 2020). Both of these observations are also consistent with the DCC dampening transcriptional variance. This “variance dampening” by the DCC, or Mof specifically, may inhibit context-dependent gene expression by reducing the ability of transcription factors to regulate expression subsequent to DCC-associated chromatin modifications that are already up-regulating expression (Table 1).

We used bacterial infection, viral infection, and abiotic stressors as model systems to test the hypothesis that the *Drosophila* DCC is a variance dampener that reduces differential expression of X-linked genes in specific contexts. Biotic and abiotic stress represents a notable contrast to previous studies of context-dependent expression involving X-autosome comparisons of genes with tissue-specific expression (e.g., Mikhaylova and Nurminsky 2011; Meisel *et al*. 2012a). Bacterial infection, for example, causes the dramatic induction of gene expression, including effectors of the humoral immune system that are expressed more than 100 times higher within 12 hours (De Gregorio *et al*. 2001; Troha *et al*. 2018; Schlamp *et al*. 2021). Curiously, none of the 30–40 *D. melanogaster* genes encoding antimicrobial peptides (AMPs, a class of effectors) are found on the X chromosome (Hill-Burns and Clark 2009), providing *a priori* evidence consistent with selection against X-linked genes induced by infection. We analyzed multiple RNA-seq studies of gene expression after biotic and abiotic treatments to test the hypothesis that the DCC inhibits context-dependent differential expression, which would explain the paucity of genes with tissue- or environment-specific expression on the *Drosophila* X chromosome.

## RESULTS

### Genes differentially expressed after infection are under-represented on the Drosophila melanogaster X chromosome

We tested if the *D. melanogaster* X chromosome is depauperate for genes induced (i.e., up-regulated) by bacterial infection regardless of functional annotation. To those ends, we analyzed RNA-seq data in which *D. melanogaster* males were infected with one of 10 different bacteria versus a control (Troha *et al*. 2018). From those infection experiments, we selected results from the five bacterial treatments with >50 differentially expressed (DE) genes, in order to have sufficient power to detect X-autosome differences. For three out of five bacterial infections we considered, there was a significant deficiency of induced genes on the X chromosome (Figure 1A). For the remaining two bacterial infections, the observed number of induced genes on the X chromosome was less than expected, although the difference was not significant. It is unlikely to observe fewer induced X-linked genes than expected for all five treatments, assuming a null hypothesis of equal proportions above and below the expectation (*p*=0.031 in a binomial exact test). In addition, when we considered all Gram-positive or Gram-negative bacteria (with a statistical model that has bacterial strain nested in treatment) from the experiment together, there was a significant deficiency of induced genes on the X chromosome in both cases (Figure 1A). Moreover, genes that were up-regulated by at least one, two, three, or four different bacteria were also significantly under-represented on the X chromosome (Figure 1B). Therefore, genes that are induced by bacterial infection are generally under-represented on the *D. melanogaster* X chromosome regardless of the criteria used to categorize induction. This is consistent with the deficiency of X-linked AMP genes (Hill-Burns and Clark 2009).

**Figure 1.**
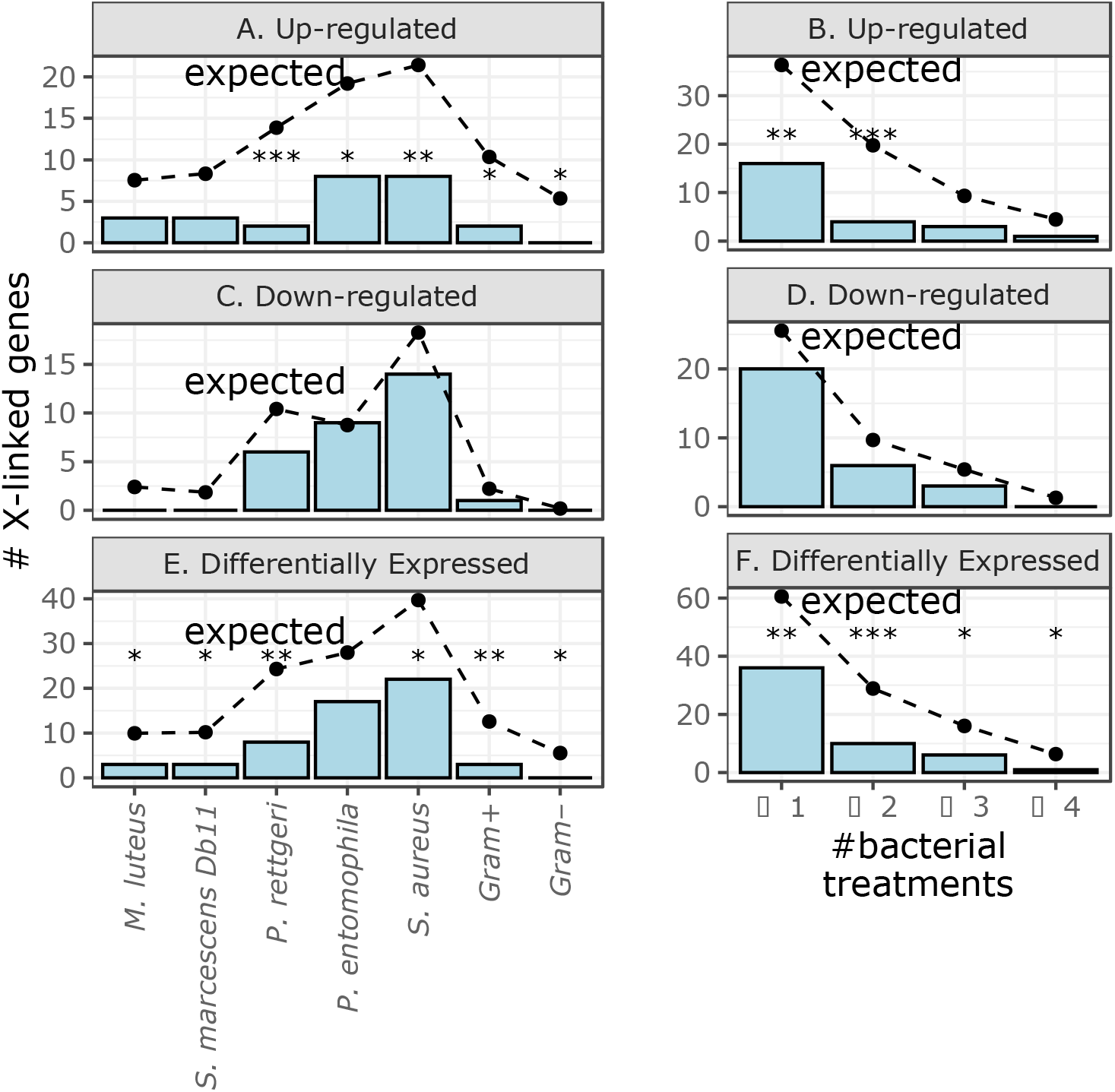
Genes that were differentially expressed after bacterial infection in *D. melanogaster* males are under-represented on the X chromosome. Bar graphs show the number of X-linked genes that were up-regulated (A-B), down-regulated (C-D), or differentially expressed (E-F) after infection with individual bacteria, combined Gram-negative (Gram–) bacteria, combined Gram-positive (Gram+) bacteria, (A, C, E), or across multiple bacterial treatments (B, D, F). Dots connected by broken lines show the expected number of X-linked genes in each category based on the fraction of autosomal genes that are differentially expressed after infection. Black asterisks represent significant differences between observed and expected counts in a Fisher’s exact test (\**P*<0.05, \**\*P*<0.005, \*\**\*P*<0.0005).

There was not a significant deficiency of X-linked *D. melanogaster* genes down-regulated after infection with any of the five bacterial treatments or when we considered all Gram-positive or Gram-negative bacteria (Figure 1C). Similarly, genes that were downregulated by one or more different bacteria were not significantly under-represented on the X chromosome (Figure 1D). However, in most cases, the number of down-regulated X-linked genes was less than the expectation, albeit not significant. The failure to detect a significant deficiency of down-regulated X-linked genes may have been caused by low statistical power— there were fewer down-regulated genes than up-regulated genes in most bacterial treatments. We examine this further by considering other experiments with more down-regulated genes below.

The total number of DE genes (either induced or repressed after infection) on the X chromosome was significantly less than the expectation for 4 of 5 individual bacterial treatments, Gram-positive bacteria, and Gram-negative bacteria (Figure 1E). The X chromosome also has significantly fewer genes that were DE in at least one or more of the bacterial treatments (Figure 1F). Therefore, both up-regulated and DE genes are under-represented on the X chromosome.

To further evaluate if the X chromosome is anomalous, we tested if any individual autosomes had a deficiency (or excess) of induced, repressed, or DE genes following bacterial infection. Notably, the left arm of the second chromosome (2L) had an excess of induced genes in every treatment (Supplemental Figures S1-S2). This is surprising because none of the annotated *D. melanogaster* AMP genes are on chromosome 2L, and only 10/74 immune effector genes are found on 2L (Sackton *et al*. 2007). Chromosome 2L therefore has an excess of genes induced by bacterial infection (Supplemental Figures S1-S2), despite having a significant deficiency of effector genes (*p*=0.006 comparing effector and non-effector immune genes on chromosome 2L with the other chromosomes in a Fisher’s exact test). The right arm of the third chromosome (3R), in contrast, has a deficiency of DE genes (Supplemental Figures S1-S2), even though it contains at least five AMP genes and has neither an excess nor a deficiency of effector genes (*p*=0.7 comparing effectors and non-effectors between 3R and other chromosomes in Fisher’s exact test). We next ranked each chromosome arm within each treatment by the percent of induced, repressed, or DE genes (excluding the diminutive chromosome 4 because it has <100 genes). On average, the X chromosome has the lowest percentage of induced, repressed, or DE genes across all treatments, and chromosome 3R has the second lowest (Supplemental Figures S3-S4). Therefore, the X chromosome is anomalous from each of the autosomes in its deficiency of up-regulated and DE genes after bacterial infection.

If the X chromosome has a maximal expression that prevents up-regulation of individual genes (i.e., a dosage limit), we should observe a difference in the distribution of log_2_FC values between X-linked and autosomal genes (Meiklejohn *et al*. 2011; Meiklejohn and Presgraves 2012). If we do not observe such a difference, it would suggest that there is not a dosage limit that selects against induced X-linked genes (Table 1). The X chromosome did not have a significantly lower log_2_FC than the autosomes in any of the five bacterial treatments (Supplemental Figure S5). In two of the five bacterial treatments (*M. luteus* and *S. marcescens Db11*), X-linked genes possess a higher median log_2_FC than autosomal genes (Supplemental Figure S5), which is opposite of the direction predicted by the dosage limit hypothesis. Therefore, the paucity of X-linked genes up-regulated after infection cannot be explained by an overall reduced dose of X chromosome gene expression.

We next tested if genes that are induced or repressed in female *D. melanogaster* following infection are also under-represented on the X chromosome. To those ends, we identified DE genes in *D. melanogaster* males and females 8 h after infection with *P. rettgeri* (Duneau *et al*. 2017), one of the bacteria that induced a deficiency of X-linked genes in males (Figure 1A). We report results for multiple log_2_FC cutoffs because raw data or *p*-values are not available for this experiment. Surprisingly, we observed a significant deficiency of X-linked genes up-regulated in males at only one of the eight log_2_FC cutoffs we considered (Figure 2A). In contrast, at five of the eight log_2_FC cutoffs, there was a significant deficiency of X-linked genes up-regulated in females after infection (Figure 2A). Similarly, at 5 of 8 log_2_FC cutoffs there was a deficiency of X-linked genes down-regulated in females (Figure 2B). Considering genes that were DE after infection, regardless of up- or down-regulation, there was a significant deficiency on the X chromosome at 7 and 4 log fold-change cutoffs in females and males, respectively (Figure 2C). Therefore, there is a significant deficiency of X-linked DE genes after infection in both male and female *D. melanogaster*.

**Figure 2.**
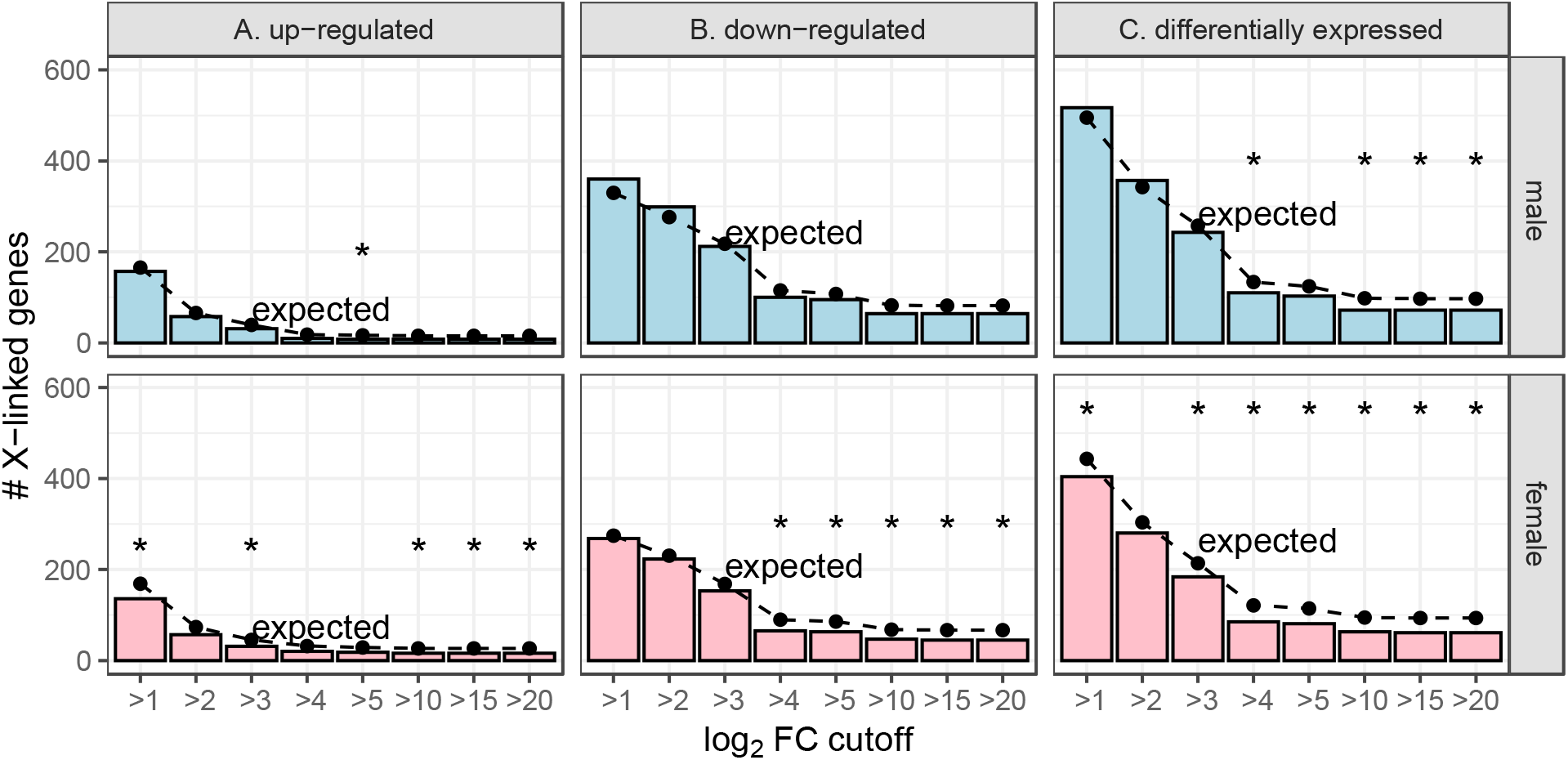
Genes that were differentially expressed after bacterial infection in *D. melanogaster* are under-represented on the X chromosome. Bar graphs show the number of X-linked genes that were up-regulated (A), down-regulated (B), or differentially expressed (C) after infection with *P. rettgeri* in either males (top) or females (bottom). Dots connected by broken lines show the expected number of X-linked genes in each category based on the fraction of autosomal genes that are differentially expressed after infection. Black asterisks represent significant differences between observed and expected counts in a Fisher’s exact test (\**P*<0.05).

We also tested if there is a paucity of X-linked genes induced at different time points following immune challenge by analyzing RNA-seq data from 1–120 h after *D. melanogaster* males were injected with *E. coli*-derived crude lipopolysaccharide (Schlamp *et al*. 2021). There was a significant deficiency of X-linked genes up-regulated at 16 of 19 time points (Figure 3A). Similarly, at 9 of 19 timepoints, there was a significant deficiency of X-linked genes downregulated (Figure 3B). Furthermore, there was a significant deficiency of X-linked DE genes (regardless of up- or down-regulation) at all time points (Figure 3C). Therefore, both up- and down-regulated genes are under-represented on the X chromosome across the full temporal spectrum during the response to infection.

**Figure 3.**
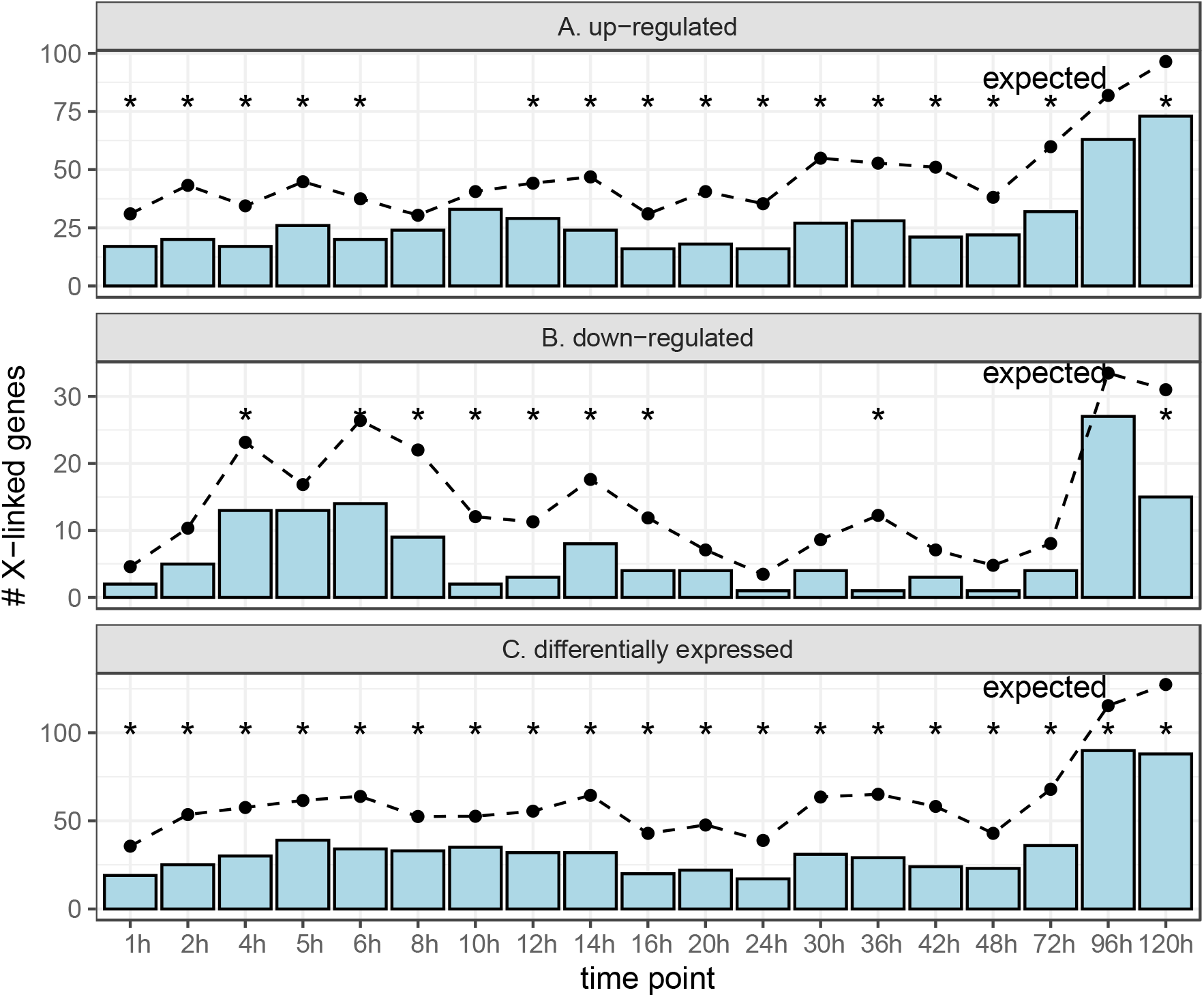
Genes that were differentially expressed at different time points after immune challenge in *D. melanogaster* males are under-represented on the X chromosome. Bar graphs show the number of X-linked genes that were up-regulated (A), down-regulated (B), or differentially expressed (C) after *D. melanogaster* males were injected with *E. coli*-derived crude lipopolysaccharide. Dots connected by broken lines show the expected number of X-linked genes in each category based on the fraction of autosomal genes that are differentially expressed after infection. Black asterisks represent significant differences between observed and expected counts in a Fisher’s exact test (\**P*<0.05).

These results show that there is a paucity of DE genes on the X chromosome after infection across a wide range of bacterial pathogens, in both sexes, and across a dense sampling of timepoints. The paucity of X-linked DE genes can be attributed to a deficiency of both up- and down-regulated genes. This suggests the deficiency of X-linked DE genes is robust to experimental variation, and also that gene dosage in males cannot fully explain the pattern.

### Genes differentially expressed under viral or abiotic stress are usually under-represented on the *Drosophila melanogaster* X chromosome

We next tested if genes induced by viral infection are under-represented on the *D. melanogaster* X chromosome. We found fewer X-linked genes than expected were induced by Zika or Kallithea virus, albeit at insignificant differences (Figure 4A). Neither viral infection resulted in a significant deviation from the expected number of X-linked down-regulated genes either (Figure 4B). The Kallithea virus data were collected from both males and females, and genes that were up-regulated in females after Kallithea infection were significantly under-represented on the X chromosome (Figure 4A). There was also a deficiency of X-linked down-regulated genes following Kallithea virus infection in males (Figure 4B). In addition, genes that were DE after Zika virus infection were under-represented on the X chromosome (Figure 4C).

**Figure 4.**
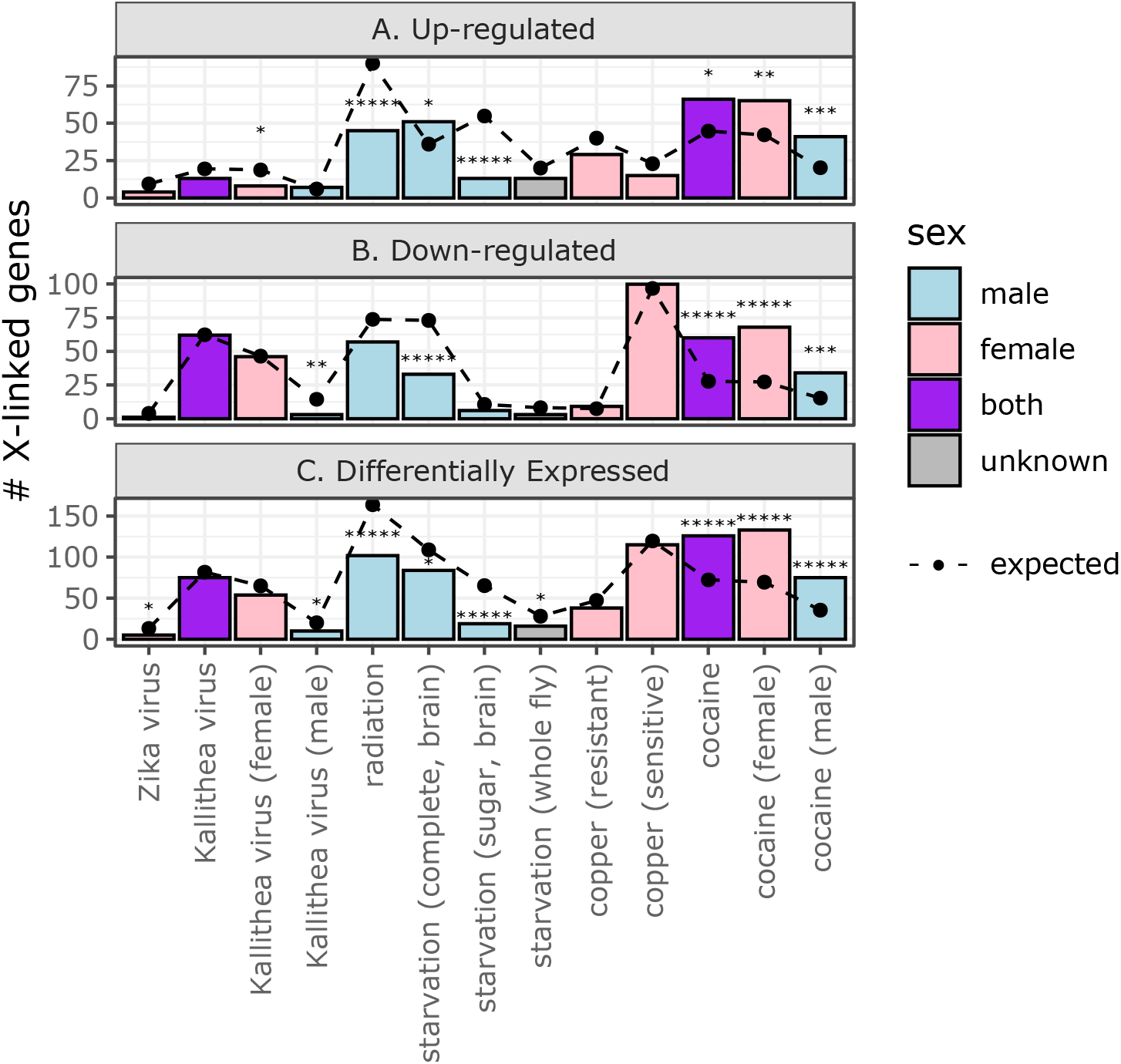
Genes that are differentially expressed after viral or abiotic treatments can be under- or over-represented on the X chromosome. Bar graphs show the number of X-linked genes that were up-regulated (A), down-regulated (B), or differentially expressed (C) after *D. melanogaster* were subjected to a viral or abiotic treatment. The treatments are listed on the x-axis, and bars are colored based on the sex of the flies used in the experiment. Dots connected by broken lines show the expected number of X-linked genes in each category based on the fraction of autosomal genes that are differentially expressed after treatment. Black asterisks represent significant differences between observed and expected counts in a Fisher’s exact test (\**P*<0.05; \*\*\*\*\**P*<0.000005).

We also tested if genes induced by four different abiotic treatments are under-represented on the X chromosome. These data include copper treatment for genotypes that are sensitive to copper and those that are resistant, which we analyzed separately. The data also included starvation treatments in which gene expression was measured in whole flies, adult brains after complete starvation, and adult brains after sugar starvation. Both radiation and sugar starvation resulted in a significant deficiency of X-linked induced genes (Figure 4A). Complete starvation (with expression measured in the brain) was the only abiotic treatment that resulted in a significant deficiency of X-linked down-regulated genes (Figure 4B). There was also a deficiency of X-linked DE genes after both radiation and starvation (Figure 4C). In contrast to all other biotic and abiotic treatments, there was an excess of X-linked genes up-regulated after complete starvation and cocaine treatment (Figure 4A). There was also an excesss of X-linked down-regulated and DE genes after exposure to cocaine, regardless of the sex of the flies (Figure 4). Both the complete starvation and cocaine treatments measured gene expression in the brain, suggesting that the brain may be an outlier with an excess, rather than a deficiency, of X-linked up-regulated genes (or DE genes in general) following abiotic stress.

We further tested if any individual autosomes had a deficiency (or excess) of DE genes following viral infection or abiotic treatments. None of the autosomal chromosome arms had a consistent excess or deficiency of induced, repressed, or DE genes after viral or abiotic treatment (Supplemental Figure S6). When we ranked each chromosome arm within each treatment by the percent of induced, repressed, or DE genes, the X chromosome had the lowest percentage of induced, repressed, or DE genes when averaged across all treatments (Supplemental Figure S7). This provides additional evidence that the X chromosome is an outlier with a deficiency of DE genes.

In summary, out of six total viral and abiotic treatments, the observed number of X-linked up-regulated genes was less than expected in most treatments, and significantly so in three treatments (Figure 4A). Down-regulated and DE genes were also under-represented on the X chromosome in some treatments (Figure 4B-C). In addition, we observed similar patterns regardless of sex or genotype of the flies used in the experiments, although sex did affect whether differences were statistically significant (Figure 4). The notable exceptions to this pattern are the effects of starvation or cocaine on gene expression in the brain, which were the only treatments (biotic or abiotic) that resulted in a significant excess of X-linked genes that were up-regulated (Figure 4).

### DE genes are less likely to be bound by the DCC

We evaluated the hypothesis that the DCC prevents the induction of X-linked genes by testing if there is a relationship between induction and DCC binding. In total, a smaller fraction of DCC-bound genes were up-regulated than X-linked unbound genes for nearly all bacterial, viral, and abiotic treatments (Figure 5A), if we do not consider whether the difference is significant. Within individual treatments, DCC-bound genes were significantly less likely to be up-regulated, relative to X-linked unbound genes, following *P. entomophila* infection, radiation treatment, starvation (in brain), copper exposure (for resistant flies), and cocaine feeding (Supplemental Figure S8A). The negative effect of DCC binding on up-regulation following cocaine was observed for both male and female flies. These results are in accordance with what would be expected if the DCC prevents induction of X-linked genes.

**Figure 5.**
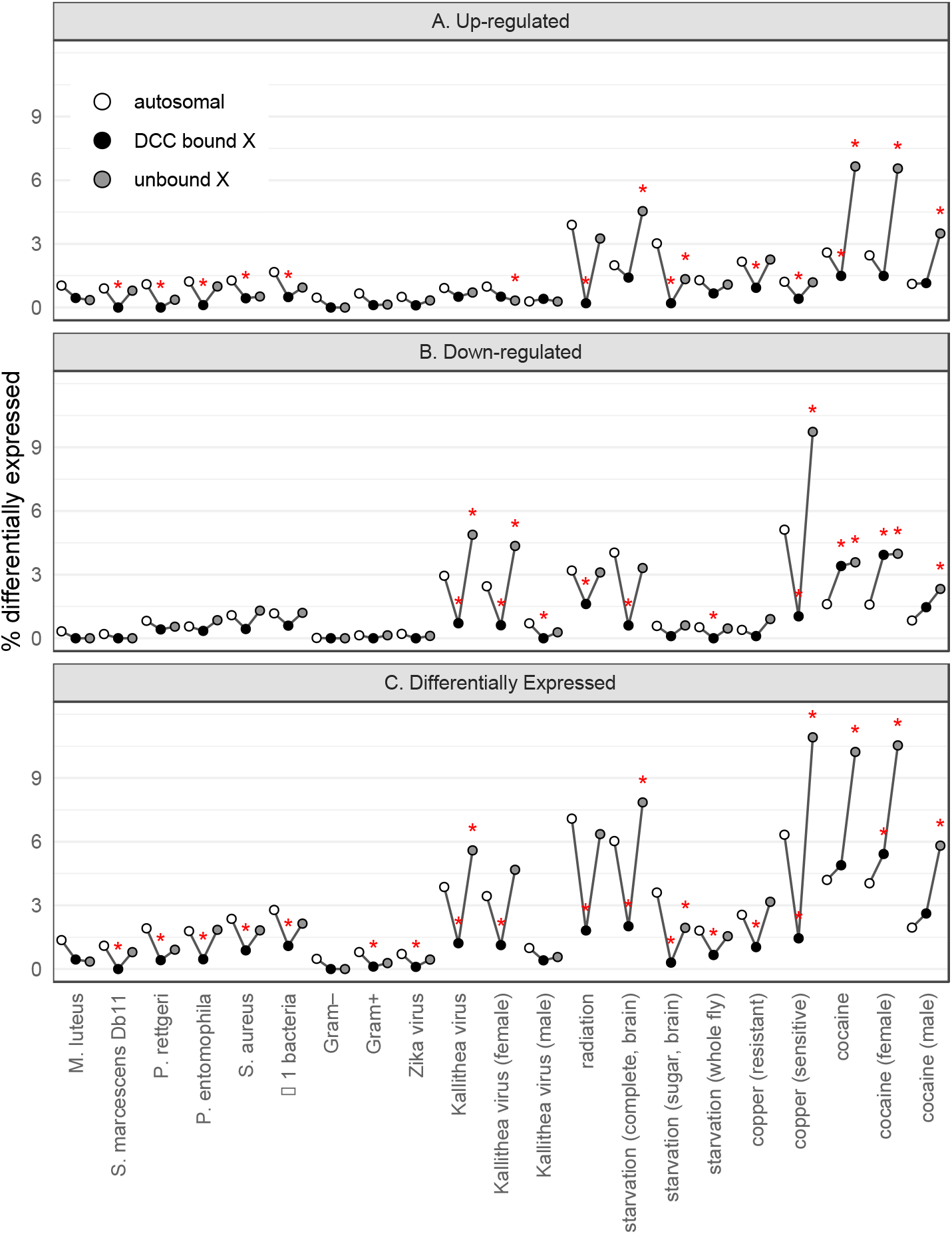
Genes that are differentially expressed (DE) after treatment are less likely to be bound by the dosage compensation complex (DCC). The percent of genes that are DE given that they are autosomal (white), X-linked and bound by the DCC (back), or X-linked and unbound (gray) is plotted for each treatment. Data are shown for genes up-regulated after a treatment (A), genes down-regulated after a treatment (B), and all DE genes (C). Red asterisks show significant differences between the number of genes that are DE and not DE when comparing autosomal vs DCC bound or unbound X-linked genes (\**P*<0.05 in Fisher’s exact test).

We next tested if DCC binding could explain the paucity of up-regulated genes on the X chromosome. To those ends, we determined if there is a difference in the proportion of up-regulated genes when we compare the autosomes with either DCC-bound or unbound genes on the X chromosome. If DCC binding explains the paucity of up-regulated genes, we expect a “V-shaped” pattern when we plot the %DE genes amongst autosomes, DCC-bound X-linked genes, and unbound X-linked genes (Figure 5). There was a significant deficiency of DCC-bound up-regulated genes relative to the autosomes across 7 different treatments (*S. marcescens* Db11, *P. rettgeri*, *P. entomophila*, *S. aureus*, radiation, copper, and cocaine), among genes up-regulated in at least one bacterial treatment, and for genes up-regulated by Gram-positive bacteria (Figure 5A). This deficiency is the bottom of the V-shape. In contrast, there was only one treatment (Kallithea virus in females) in which there was a significant deficiency of up-regulated X-linked genes unbound by the DCC relative to autosomal genes (Figure 5A). Most other treatments had the V-shape, with no significant differences in the fraction of DE genes between autosomal and unbound X-linked genes. In addition, unbound X-linked genes were significantly more likely to be up-regulated by cocaine than autosomal genes (Figure 5A). Therefore, the evidence for a paucity of X-linked up-regulated genes is much greater for DCC-bound than unbound genes (creating the V-shape in Figure 5A), which is consistent with the expectation if the DCC prevents the induction of X-linked genes.

We also observed that X-linked genes bound by the DCC were less likely to be down-regulated after treatment than X-linked unbound genes (Figure 5B). In three different viral or abiotic treatments (Kallithea virus, radiation, and copper), X-linked down-regulated genes were significantly less likely to be DCC-bound than unbound (Supplemental Figure S8B). The same results were observed for Kallithea viruses when we considered female samples only, and similar trends were observed for males (although not significant because of small sample sizes of down-regulated genes). Therefore, the DCC appears to interfere with both up- and down-regulation of gene expression. The cumulative effect of the DCC on both up- and down-regulation can be seen in the significant deficiency of DCC-bound DE genes for five unique treatments (Supplemental Figure S8C).

DCC-bound genes were also significantly less likely to be down-regulated than autosomal genes in four different treatments—Kallithea virus, radiation, starvation, and copper (Figure 5B). Similarly, DCC-bound genes were less likely to be DE than autosomal genes after most treatments (Figure 5C). In contrast, X-linked unbound genes were significantly more likely to be down-regulated or DE than autosomal genes after Kallithea virus infection, copper treatment, or cocaine (Figure 5). Therefore, down-regulated and DE genes also tend to have the V-shaped distribution. These results are all consistent with the expectations if the DCC prevents down-regulation of X-linked genes.

Our results provide consistent evidence that DCC binding can largely explain the paucity of X-linked up-regulated, down-regulated, and DE genes. Notably, we observed much stronger evidence for a deficiency of X-linked DE genes when we considered DCC-bound genes, and only weak (or no) evidence for unbound genes (the V-shapes in Figure 5). This is consistent with the hypothesis that the DCC is a variance dampener that prevents both up- and down-regulation of X-linked genes. However, there is a deficiency of up-regulated X-linked unbound genes relative to autosomal genes in some treatments (Figure 5A), suggesting that DCC binding alone cannot completely explain the paucity of up-regulated genes on the X chromosome. Therefore, other factors, such as a dosage limit, may also be necessary to explain the exclusion of up-regulated genes from the *Drosophila* X chromosome.

### Differentially expressed genes are further from DCC high affinity sites

A complementary way to assess the effect of the DCC on differential gene expression is to measure the distance to the nearest DCC high affinity site (HAS) for each gene. We cannot test for differences in distance to HAS between DE genes and non-DE genes because there are too few X-linked DE genes for statistical testing. Instead, we calculated the correlation between distance to the nearest HAS and log_2_ fold-change between treatment and control (log_2_FC) for all genes, regardless of whether they are significantly DE.

First, we considered |log_2_FC| as a measure of the extent of differential expression, regardless of up- or down-regulation. In nearly all combinations of treatments, HAS data sets, and sexes, there was a positive correlation between |log_2_FC| and distance from an HAS (Figure 6A). Therefore, X-linked genes further from an HAS were more differentially expressed following bacterial infection, viral infection, or abiotic treatment. This is consistent with the hypothesis that the DCC inhibits differential expression (i.e., both up- and down-regulation).

**Figure 6.**
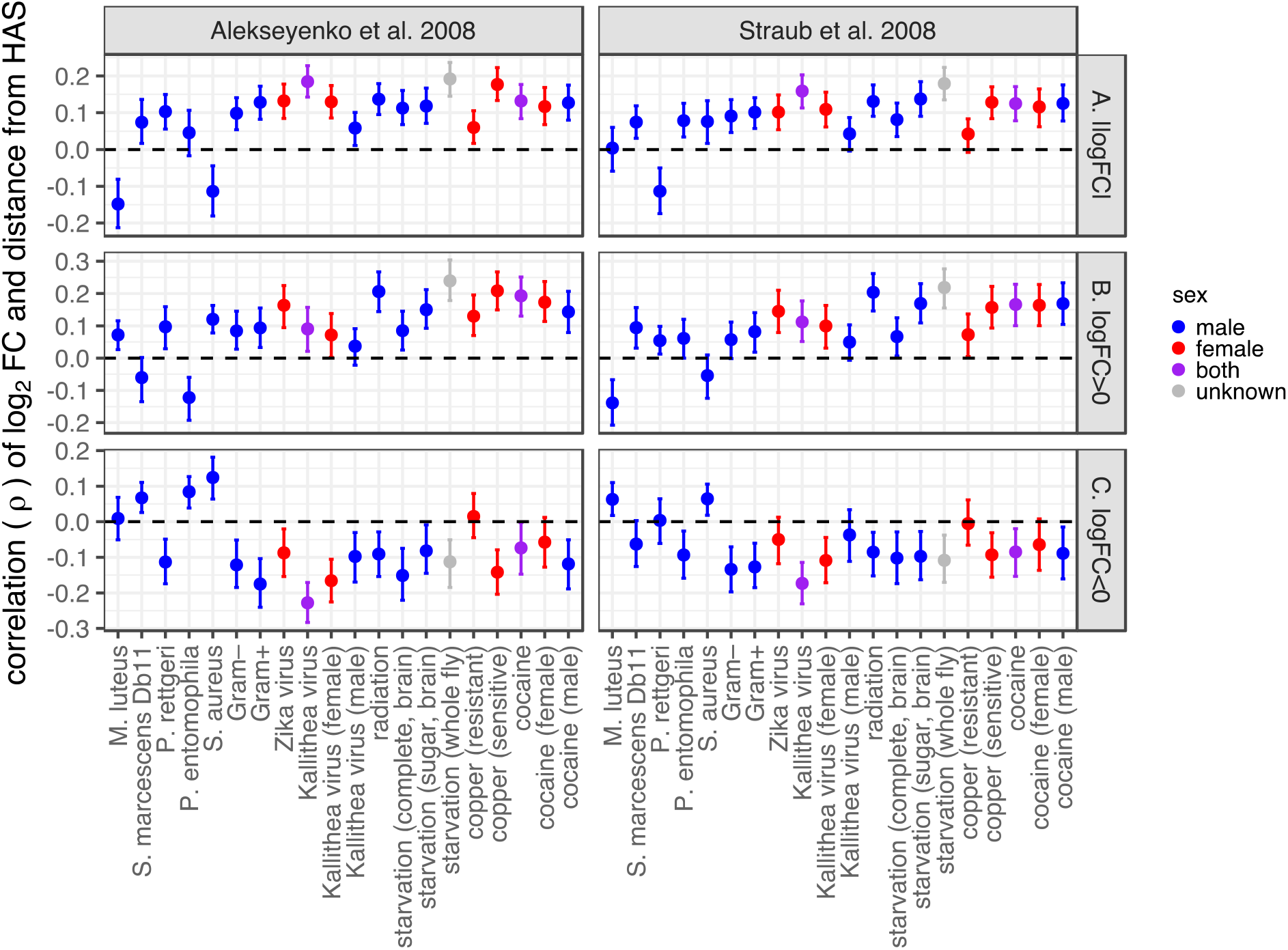
Correlations between distance from a dosage compensation complex high affinity site (HAS) and the log_2_ fold-change in expression between treatment and control (log_2_FC). Each dot is the rank order correlation (ρ) between distance to the nearest HAS and log_2_FC. Error bars show the 95% confidence interval determined by bootstrap resampling the data 1,000 times. The X-axis shows the specific treatment. Dots and error bars are colored based on the sex of the flies used in the experiment (see legend). HAS were obtained from two different data sets (Alekseyenko *et al*. 2008; Straub *et al*. 2008), with results from the two different data sets shown separately in the two columns. Correlations are plotted with |log_2_FC| values for all genes (A), only genes with log_2_FC>0 (B), and only genes with log_2_FC<0 (C).

To specifically evaluate if proximity to an HAS affects up-regulation, down-regulation, or both, we separately considered genes with log_2_FC>0 and log_2_FC<0 (regardless of whether the deviation from 0 is significant). When we considered only genes with log_2_FC>0, there was evidence for a positive correlation between log_2_FC and distance from an HAS for most treatments (Figure 6B). In comparison, when we considered genes with log_2_FC<0, there was a negative correlation between log_2_FC and distance from an HAS for most treatments (Figure 6C). Both of these correlations indicate that genes further from an HAS were more likely to be either up- or down-regulated after bacterial infection, viral infection, or abiotic treatment. These results are consistent with the hypothesis that the DCC inhibits both up- and down-regulation of gene expression.

### Differential expression, dosage compensation, and expression breadth

We next considered if expression breadth could explain the correlations between log_2_FC and distance from an HAS. This analysis was motivated by the previously described observation that genes bound by the DCC or closer to an HAS are narrowly expressed in fewer tissues than X-linked unbound genes and those further from an HAS (Meisel *et al*. 2012b). Here, we calculated partial correlations (Schäfer and Strimmer 2005) between log_2_FC, distance from an HAS, and expression breadth. We quantified expression breadth using τ, which ranges from 0 (for genes expressed in many tissues) to 1 (for genes highly expressed in a single tissue) (Yanai *et al*. 2005). We confirmed the positive correlation between τ and distance from an HAS, even when log_2_FC is included in the analysis (Supplemental Figures S9–S12). We also found that |log_2_FC| was positively correlated with τ (Supplemental Figures S9–S12). Moreover, there was a positive correlation between log_2_FC and τ for genes with log_2_FC>0, and there was a negative correlation for genes with log_2_FC<0 (Supplemental Figures S9–S12). Therefore, genes that were more up- or down-regulated tended to also be more narrowly expressed. Unsurprisingly, genes induced by bacterial infection were narrowly expressed in the fat body, which is the primary organ of the humoral immune response (Lemaitre and Hoffmann 2007).

When we considered the correlation with τ, many partial correlations between log_2_FC and distance from an HAS were no longer significantly different from zero (Figure 7; Supplemental Figure S13). This is true regardless of whether sex-specific reproductive tissues are included in the calculation of τ. Therefore, the correlations between log_2_FC and distance from an HAS could often be explained by the correlations between τ and both log_2_FC and distance from an HAS. However, some partial correlations between log_2_FC and distance from an HAS still remained significantly different from 0 (Figure 7). These significant correlations were almost always in a direction consistent with genes further from an HAS being more up-regulated or more down-regulated (i.e., a positive correlation for |log_2_FC| or log_2_FC>0, or a negative correlation for log_2_FC<0). The one exception to this rule was a positive partial correlation between log_2_FC and distance from an HAS for genes with log_2_FC<0 after copper treatment in resistant flies (Figure 7D). This negative correlation is suggestive that genes closer to an HAS are more down-regulated following copper treatment. However, there were very few genes down-regulated after copper treatment in resistant flies (Figures 4–5), suggesting that this positive correlation may be an artifact of a small range of negative log_2_FC values.

**Figure 7.**
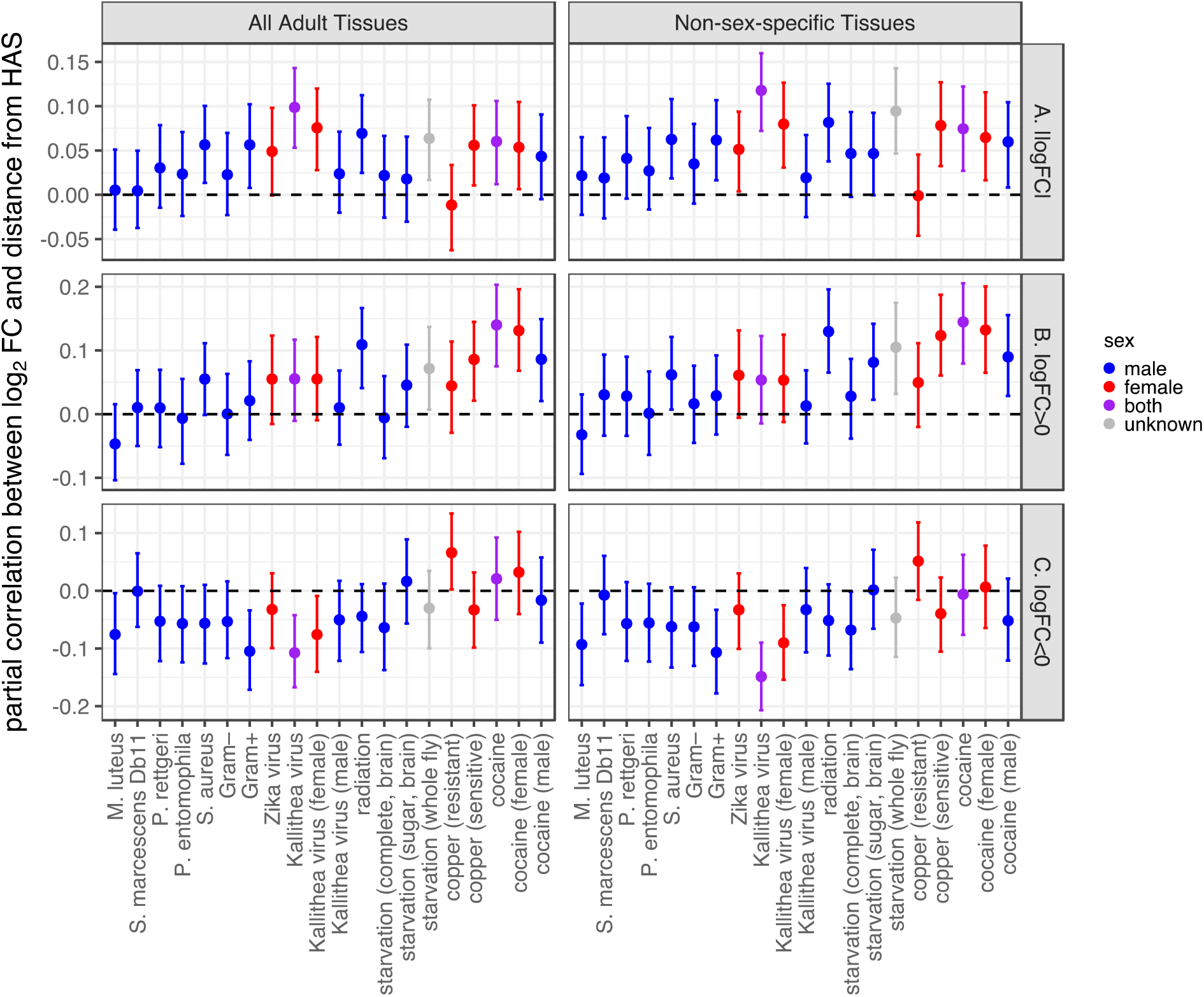
Partial correlations between log_2_ fold-change in expression between treatment and control (log_2_FC) and distance from a dosage compensation complex high affinity site (HAS). Partial correlations were calculated based on rank order correlations between log_2_FC, distance from an HAS, and tissue expression breadth. Each dot shows the partial correlation between log_2_FC and distance from an HAS, with the error bars representing 95% confidence intervals from 1,000 bootstrap replicates of the data. The X-axis shows the specific treatment. Dots and error bars are colored based on the sex of the flies used in the experiment (see legend). HAS were obtained from the Alekseyenko *et al*. (2008) data set; results from the Straub *et al*. (2008) data are shown in the supplemental material. Expression breadth was calculated using microarray data from either 14 unique adult tissues (left) or 10 adult tissues that are not sex-specific (right). Partial correlations are plotted with log_2_FC values for all genes (A), |log_2_FC| values for all genes (B), only genes with log_2_FC>0 (C), and only genes with log_2_FC<0 (D).

## DISCUSSION

We analyzed RNA-seq data from *D. melanogaster* subjected to bacterial, viral, or abiotic treatments. We found that DE genes—both up- and down-regulated—were often depauperate on the X chromosome regardless of the sex of the flies or the type of treatment (Figures 1–4).

We additionally determined that DE genes were less likely to be bound by the DCC, and that DCC binding can largely explain the deficiency of X-linked DE genes (Figure 5). In addition, genes that are further from an HAS were more differentially expressed, regardless of whether the genes were up- or down-regulated (Figure 6). However, much of the relationship between differential expression and distance from an HAS could be explained by both variables being correlated with tissue-specific gene expression (Supplemental Figures S9–S12). Nonetheless, a significant correlation between log_2_FC and distance from an HAS remained for some treatments after controlling for expression breadth across tissues (Figure 7).

### Chromosome 2L has a complementary gene content to the X chromosome

An excess of chromosome 2L genes were up-regulated in many conditions, in contrast to the deficiency of X-linked DE genes. We found an excess of chromosome 2L genes induced in every bacterial treatment (Supplemental Figures S1-S4), even though no AMP genes (and only 10 of 74 immune effectors) are on chromosome 2L (Sackton *et al*. 2007). An excess of chromosome 2L genes were also induced after copper treatment (Supplementary Figure S6). Previous analyses revealed that chromosome 2L (also known as Muller element B) has an excess of genes with male-biased expression in multiple *Drosophila* species, in contrast to the paucity of X-linked genes up-regulated in males (Parisi *et al*. 2003; Meisel *et al*. 2012a). In addition, chromosome 2L has an excess of genes encoding accessory gland proteins, while the X chromosome is depauperate (Ravi Ram and Wolfner 2007). The expression level in accessory gland is also higher for chromosome 2L genes, and lower for X-linked genes, than other chromosomes (Meisel *et al*. 2012a). Moreover, chromosome 2L genes have narrower expression in specific tissues than genes on other chromosomes, in contrast to X-linked genes that tend to be more broadly expressed (Meisel *et al*. 2012a). Future work could address why chromosome 2L has a complementary pattern to the X chromosome.

### Minimal support for the dosage limit hypothesis

Our analysis allowed us to test the dosage limit hypothesis, which predicts that highly expressed genes (i.e., those that are up-regulated in specific contexts) are under-represented on the X chromosome because of its haploid dose in males (Vicoso and Charlesworth 2009; Meisel *et al*. 2012a; Hurst *et al*. 2015). We determined that *D. melanogaster* genes that are either up- or down-regulated following biotic or abiotic treatments are frequently under-represented on the X chromosome (Figures 1–4). The paucity of down-regulated X-linked genes is inconsistent with the dosage limit hypothesis, which only predicts that up-regulated genes will be under-represented on the X chromosome (Table 1). Moreover, we frequently observed a deficiency of up-regulated X-linked genes in females, but not males, when data from both sexes were available (Figures 2 and 4). A female-specific paucity of X-linked up-regulated genes is also not predicted by the dosage limit hypothesis. Furthermore, there was an excess of X-linked up-regulated genes following cocaine treatment in both males and females (Figure 4), which is also inconsistent with the dosage limit hypothesis.

Our results thus provide strong evidence that the dosage limit hypothesis cannot completely explain the unique gene content of the *Drosophila* X chromosome. However, we also cannot explain the paucity of X-linked up-regulated genes based on proximity to an HAS alone (Figure 5). A dosage limit may therefore be partially responsible for some aspects of the unique gene content of the X chromosome. The dosage limit hypothesis may be especially important in testis, where the haploid dose of the X chromosome does not appear to be compensated (Meiklejohn *et al*. 2011). Specifically, the reduced dosage of the X chromosome could explain the biased duplication of genes from the X to the autosomes, with the autosomal derived paralogs expressed primarily in testis in order to compensate for under-expression of the X (Betrán *et al*. 2002; Meisel *et al*. 2009, 2010). In addition, the dosage limit hypothesis is also consistent with the observation that genes expressed specifically in the male accessory gland are under-represented on the X chromosome (Swanson *et al*. 2001; Ravi Ram and Wolfner 2007; Meisel *et al*. 2012a). Therefore, both dosage limits and DCC binding may act in concert to exclude up-regulated genes from the *Drosophila* X chromosome.

### Evidence for the DCC as a variance dampener that inhibits X-linked DE genes

Our results add evidence in support of the hypothesis that the DCC is a variance dampener (Lee *et al*. 2018), which prevents both up- and down-regulation of gene expression on the *Drosophila* X chromosome. First, there is a significant deficiency of down-regulated genes on the X chromosome, in addition to a paucity of X-linked up-regulated genes, in a variety of treatments (Figures 2–4). Second, the X-linked genes that are up- or down-regulated tend not to be bound by the DCC, and DCC-binding largely explains the X-autosome differences in up-regulated, down-regulated, and DE genes (Figure 5). Lastly, the magnitude of differential expression is correlated with distance from an HAS for both up- and down-regulated genes (Figures 6 and 7). Because we observe similar results for both up- and down-regulated genes, we hypothesize that the DCC is a variance dampener that inhibits both induction and repression of gene expression (Table 1).

We further hypothesize that the variance dampening effect of the DCC can explain other aspects of X chromosome gene content. For example, there is a paucity of genes with tissue-specific expression on the *Drosophila* X chromosome (Mikhaylova and Nurminsky 2011; Meisel *et al*. 2012a). The genes with tissue-specific expression that are on the X chromosome are less likely to be DCC-bound than broadly expressed X-linked genes (Meisel *et al*. 2012b). Tissue-specific expression can be thought of as a form of context-dependence in which the context is developmental, rather than environmental. Therefore, the variance dampening effect of the DCC may prevent tissue-specific induction, or repression, of gene expression.

There are at least two explanations for how the variance dampening effect of the DCC could lead to a deficiency of genes with context-specific expression on the X chromosome. First, in what we will call the mechanistic explanation, the DCC itself may affect gene expression in the experiments from which we obtained data. While this mechanism is feasible for gene expression in males, it is less obvious how it could explain the deficiency of X-linked DE genes in females (Figures 2 and 4) because the DCC does not assemble in females (Lucchesi and Kuroda 2015). The lack of a complete DCC in females occurs because the Msl-2 protein is only expressed in males (Kelley *et al*. 1997). However, Mof, the component of the DCC responsible for H4K16ac, is expressed in both sexes, and there is evidence that it serves as a variance dampener in both males and females (Lee *et al*. 2018). In addition, the *Drosophila* X chromosome has a distinct chromatin environment from the autosomes in females, including histone marks associated with dosage compensation (Zhang and Oliver 2010). These chromatin modifications, possibly mediated by Mof, may serve to dampen variance in X-linked gene expression in both sexes, and therefore reduce differential expression on the X chromosome. Furthermore, there is a complex interplay between heterochromatin and dosage compensation in *Drosophila (Makki and Meller 2021),* which could also contribute to a variance dampening effect in both sexes.

Alternatively, in what we call the evolutionary explanation, we may have observed an evolved difference between the X and autosomes in the experimental data we analyzed. These evolved differences would be a result of selection against genes with context-dependent gene expression on the X chromosome as a result of the variance dampening effect of the DCC, or perhaps only Mof. This explanation is attractive because it allows for selection in males only (where the DCC assembles) to shape the gene expression profile of the X chromosome in both sexes, where we observe the paucity of X-linked genes with context-dependent expression. This is analogous to how a shared genome prevents the evolution of sexual dimorphism because of inter-sexual phenotypic correlations (Rowe *et al*. 2018). Consistent with the evolutionary explanation, DCC bound genes have slower evolving gene expression levels than X-linked unbound genes (Meisel *et al*. 2012b), suggesting that the DCC may inhibit the evolution of gene expression. Similarly, X-linked genes whose expression levels changed more during a laboratory evolution experiment were further from an HAS (Abbott *et al*. 2020). It is worth noting, however, that the mechanistic and evolutionary explanations are not mutually exclusive, and additional work is necessary to evaluate how well each can explain the effect of the DCC, or Mof in particular, on the unique gene content of the *Drosophila* X chromosome.

Regardless of the mode of action, the variance dampening hypothesis could also explain the differences in gene content between mammalian and *Drosophila* X chromosomes. The mammalian X has an excess of genes with tissue-specific expression (Lercher *et al*. 2003; Meisel *et al*. 2012a), in contrast to the deficiency of genes with context-dependent expression on the *D. melanogaster* X chromosome. A dosage limit hypothesis has been proposed to explain the excess of narrowly expressed genes on the mammalian X chromosome because broadly expressed genes have a higher maximal expression (Hurst *et al*. 2015). Evidence for large-scale up-regulation of the mammalian X is much weaker than the evidence that the *Drosophila* DCC up-regulates X-linked expression (Xiong *et al*. 2010; Pessia *et al*. 2012; Lin *et al*. 2012; Deng *et al*. 2013; Mank 2013; Gu and Walters 2017; Deng and Disteche 2019). Therefore, the taxon-specific peculiarities of dosage compensation could explain the differences in X chromosome gene content between *Drosophila* and mammals. Specifically, only in *Drosophila* is there evidence for a variance dampener that targets X-linked genes, whereas dosage limits may be more pronounced in mammals.

### Exceptions to the rules are also evidence for the DCC as a variance dampener

We observed exceptions to the general relationships between DE genes, X-linkage, and the DCC within specific treatments. For example, there was an excess of DE genes on the X chromosome after cocaine or starvation treatments (Figure 4). In addition, some of the correlations between log_2_FC and distance from an HAS were in opposite directions depending on the treatment (Figures 6 and 7). We discuss these exceptions below, explain how they are consistent with the variance dampener hypothesis despite the atypical patterns, and describe how they can help us discriminate between the mechanistic and evolutionary explanations for the reduced DE on the X chromosome.

#### Excess of X-linked DE genes in brain

In both of the brain samples analyzed (starvation and cocaine treatments), we observed an excess of X-linked DE genes (Figure 4). The cocaine treatment resulted in an excess of both up- and down-regulated genes, whereas the starvation treatment only caused an excess of up-regulated genes. There is a similar enrichment of X-linked genes with male-biased expression in *D. melanogaster* brain (Huylmans and Parsch 2015). Multiple genes that encode DCC proteins, including *msl-2* and *mle*, are highly expressed in brain (Chintapalli *et al*. 2007; Straub *et al*. 2013; Vensko and Stone 2014; Huylmans and Parsch 2015), as are the non-coding RNAs that assemble with the DCC (Amrein and Axel 1997; Meller *et al*. 1997). If the DCC inhibits context-dependent differential expression, we may expect fewer X-linked DE genes in the brain because of the high expression of the DCC components. It is therefore surprising that we observe the opposite pattern in the brain.

Despite the excess of X-linked up-regulated genes in the brain after either starvation or cocaine, we observed that DE genes in both treatments are less likely to be bound by the DCC, consistent with most other treatments (Figure 5). Moreover, X-linked DE genes after cocaine treatment were further from an HAS, similar to other treatments (Figure 7). These observations are consistent with the hypothesis that the DCC is a variance dampener, but they do not explain why the unbound X-linked genes are more likely to be DE in the brain than in other tissues.

One explanation for an excess of X-linked DE genes in the brain is the heterogeneity of dosage compensation across the X chromosome in brain cells. Belyi et al. (2020) observed greater position effects on brain gene expression than expression in non-brain head tissus for transgenes inserted on the X chromosome. This suggests that the X chromosome is more of a patchwork of DCC-bound and unbound regions in the brain than in other tissues. The unbound regions may allow for context-dependent transcriptional regulation of X-linked genes in the brain more than in other tissues where DCC-binding is more uniform.

#### Excess of X-linked DE genes after cocaine treatment

The excess of X-linked DE genes after cocaine treatment points to a possible mechanism to explain the context-dependent transcriptional regulation of X-linked genes unbound by the DCC. Cocaine affects chromatin in neural cells by reversing trimethylation of histone H3 at lysine 9 (H3K9me3), a repressive chromatin mark, leading to de-repression of genes and repetitive elements that would normally be silenced (Kumar *et al*. 2005; Renthal *et al*. 2007; Maze *et al*. 2011; Covington *et al*. 2011). We observed that the majority of X-linked genes that were up-regulated following cocaine treatment in *D. melanogaster* brain were not bound by the DCC (Figure 5A), and previous work showed that unbound genes are more likely to be associated with repressive chromatin marks than DCC-bound genes (Meisel *et al*. 2012b). Therefore, the X-linked DE genes after cocaine treatment are probably located in chromatin that was unbound by the DCC, and cocaine induced a conversion from repressive to transcriptionally active chromatin. Importantly, this interpretation is consistent with the hypothesis that the DCC is a variance dampener and X-linked genes unbound by the DCC are more likely to have context-dependent expression. Future work should evaluate if chromatin state was in fact altered in X-linked unbound genes in neuronal cells following cocaine treatment.

#### Genotypic differences help discriminate between mechanistic and evolutionary explanations

We observed different patterns following copper treatment depending on whether the flies were sensitive or resistant. Many more X-linked genes were down-regulated in sensitive flies, and this was accompanied by an equivalent enrichment of down-regulated autosomal genes (Figure 4B). The down-regulated X-linked genes were extremely biased toward those unbound by the DCC (Figure 5B), consistent with our hypothesis that the DCC is a variance dampener. However, there was a difference between sensitive and resistant flies in the correlation of log_2_FC and distance from an HAS. In sensitive flies, the correlation was positive when we considered |log_2_FC| or genes with log_2_FC>0 (Figure 7A–B), consistent with more differential expression further from an HAS. In resistant flies, the correlation was positive for genes with log_2_FC<0 (Figure 7C), suggesting that genes further from an HAS were less down-regulated. One possible explanation for this discrepancy is that, because there were so few DE genes in resistant flies, the correlation between log_2_FC and distance from an HAS differed from other treatments (possibly because few genes have extreme negative log_2_FC values). This explanation is not well supported because other treatments had even fewer DE genes (e.g., Zika virus and starvation), yet the correlations between log_2_FC and distance from an HAS in those other treatments were in the same directions as in the treatments with more DE genes (Figures 6 and 7). Alternatively, resistance to copper (or any treatment) may affect the relationship between gene expression and the DCC.

Regardless of the causes of the differences between copper sensitive and resistant flies, the comparison is informative for evaluating the mechanistic and evolutionary explanations for the relationship between DE genes and DCC binding. Copper resistant flies are likely to resemble genotypes that would emerge following adaptation to copper exposure. As described above, X-linked DE genes in copper resistant flies were not necessarily further from an HAS (Figure 7). In contrast, X-linked DE genes in copper sensitive flies were further from an HAS, similar to what was observed in most other treatments (Figure 7). Therefore, adaptation may not necessarily lead to the evolved relationship between X-linked DE genes and the DCC that we observed, which could be interpreted as support for the mechanistic explanation for X-autosome differences. However, there are clear evolved differences between the X and autosomes, such as the absence of X-linked AMP genes (Hill-Burns and Clark 2009) and deficiency of X-linked accessory gland expressed genes (Swanson *et al*. 2001; Ravi Ram and Wolfner 2007; Meisel *et al*. 2012a), which cannot be explained by the mechanistic explanation. This leads us to conclude that both mechanistic effects of the DCC (or Mof) and evolved X-autosome differences are responsible for the paucity of X-linked DE genes. The evolved X-autosome differences are likely the result of selection against X-linked genes with context-dependent expression in response to both the variance dampening effects of the DCC/Mof and dosage limits of the haploid X chromosome in males.

## MATERIALS AND METHODS

### General statistical analysis

The analyses and figure generation were performed in R (R Core Team 2019) using the following packages: boot (Davison and Hinkley 1997; Canty and Ripley 2021), corpcor (Schäfer and Strimmer 2005; Schafer *et al*. 2017), ggplot2 (Wickham 2016), ggridges (Wilke 2021), and cowplot (Wilke 2020). Additional R packages were used for specific analyses, as described below.

### RNA-seq data analysis

We analyzed available RNA-seq data to test for differential expression between control *D. melanogaster* and flies that received a bacterial, viral, or abiotic treatment (Supplementary File 1). In one dataset, *D. melanogaster* adult males were infected with one of ten different bacteria or a control treatment (Troha *et al*. 2018). For that experiment, we used RNA-seq data from 12 hours post treatment for only live (not heat-killed) bacteria, which we downloaded as fastq files from the NCBI sequence read archive (BioProject PRJNA428174). We only included bacterial treatments with at least 50 DE genes relative to the control condition (see below for methods used to identify DE genes), which was 5/10 treatments. In another data set, we compared gene expression 8 h following injection of *Providencia rettgeri* with uninfected flies, in males and females separately (Duneau *et al*. 2017). In a third data set, we analyzed the response to infection at 19 different timepoints from 1 h to 120 h after injection of *E. coli*-derived crude lipopolysaccharide (Schlamp *et al*. 2021). Other datasets include exposure to one of two different viral infections, copper, starvation, radiation, and cocaine (Palmer *et al*. 2018; Prasad and Hens 2018; Harsh *et al*. 2020; de Oliveira *et al*. 2021; Green *et al*. 2021; Baker *et al*. 2021). A full list of accession numbers is provided in Supplementary File 1.

Raw RNA-seq reads were assigned to annotated *D. melanogaster* transcripts (r6.22) using kallisto v0.44.0 (Bray *et al*. 2016). Read counts for all transcripts from each gene were summed to obtain gene-level expression estimates, and the counts per gene were then rounded to the nearest integer. For a given treatment, we only considered genes with at least 10 mapped reads total across all replicates from control and treatment samples. The integer counts were used as input into DESeq2 (Love *et al*. 2014), which we used to identify DE genes between the treatment and control samples (see below). We performed a principal component analysis (PCA) on regularized log transformed read counts to identify replicate samples that were outliers relative to other replicates of the same sample type. We identified one outlier male control replicate in the Kallithea virus data, which we excluded from all subsequent analyses.

We identified DE genes based on the false-discovery rate (FDR) corrected p-value (p_ADJ_) and log_2_FC of treatment to control expression. Genes were considered up-regulated in a treatment if p_ADJ_ < 0.05 and log_2_FC > 1 (i.e., a significant 2-fold increase upon treatment). Similarly, genes were considered downregulated if p_ADJ_ < 0.05 and log_2_FC < -1 (i.e., a significant 2-fold decrease upon treatment). DE genes are those that are either up- or down-regulated in a given treatment (i.e., p_ADJ_ < 0.05 and |log_2_FC| > 1). We only included datasets with at least 50 DE genes between treatment and control samples.

We also analyzed the bacterial infection data considering all Gram-negative or Gram-positive bacteria from Troha *et al*. (2018) as a treatment group. In this case, we used a statistical model that examined the effect of treatment (bacteria or control), with strain nested within bacterial treatment. This was done separately for all Gram-negative bacteria and all Gram-positive bacteria. DE genes, as well as up- and down-regulated genes, were identified using the same p_ADJ_ and log_2_FC criteria described above.

In cases where both female and male RNA-seq data were available (cocaine and Kallithea virus), we performed two separate analyses focusing on the effect of the treatment (i.e., not on the effect of sex) on gene expression. First, we analyzed the male and female data together using a linear model that included the effect of treatment and the interaction of treatment and sex. From this analysis, we extracted genes that were DE based on the treatment effect. Second, we analyzed data from the two sexes separately using a model that only included the effect of treatment (i.e., the same way we analyzed data from other treatments without separate sex samples).

We did not analyze raw RNA-seq data (i.e., Illumina sequence reads) for two of the infection datasets. First, we obtained previously identified DE genes from the time course analysis of expression following infection (Schlamp *et al*. 2021). Second, raw data were not available from an experiment in which male and female *D. melanogaster* were infected with bacteria (Duneau *et al*. 2017), but processed data were available from FlySexsick-*seq* (http://flysexsick.buchonlab.com). For those data, we compared gene expression between unchallenged flies and 8 h following injection of *Providencia rettgeri*. Only expression levels (and no p-values) were provided for these data, and we therefore considered genes to be differentially expressed based on a variety of log_2_FC cutoffs.

We determined a null expectation for the number of X-linked DE genes by multiplying the fraction of autosomal genes that are DE by the total number of X-linked genes with expression measurements. Similar calculations were performed to determine a null expectation for up- or down-regulated X-linked genes.

### DCC binding

Data on DCC binding in the *D. melanogaster* genome was obtained from a published chromatin immunoprecipitation followed by microarray (ChIP-chip) experiment in which genes were classified as bound by the DCC in SL2 embryonic cells, clone 8 wing imaginal disc cells, and embryos (Alekseyenko *et al*. 2006). For the purpose of our analysis, we considered a gene to be bound by the DCC if it was bound in at least one of the three samples. We also obtained HAS locations from two different published data sets (Alekseyenko *et al*. 2008; Straub *et al*. 2008). Using these HAS locations, we calculated the distance of each X chromosome gene to the nearest HAS in nucleotides.

### Tissue-specific expression

We obtained microarray measurements of gene expression from 14 unique adult *D. melanogaster* tissues from FlyAtlas (Chintapalli *et al*. 2007). Of the 14 tissues, 4 are sex-specific (testis and accessory gland from males, and ovary and spermatheca from females), and the remaining 10 tissues (brain, eye, thoracicoabdominal ganglion, salivary gland, crop, midgut, malpighian tubule, hindgut, heart, and fat body) are shared by both males and females (i.e., non-sex-specific). We used these data to calculate expression breadth (τ) for each gene:

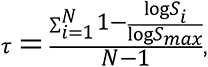

where *N* is the number of tissues analyzed (10 for the non-sex-specific tissues, and 14 for all adult tissues), *S_i_* is the gene expression level in tissue *i* (measured as average signal intensity for all microarray probes assigned to that gene), and *S_max_* is the maximum *S_i_* across all *N* tissues (Yanai *et al*. 2005). All *S_i_* < 1 were set to 1 so that log *S_i_* ≥ 0, as done previously (Larracuente *et al*. 2008; Meisel 2011; Meisel *et al*. 2012a). Values for spermatheca from mated and unmated females were averaged to create a single *S_i_* for spermatheca (Meisel 2011). We calculated τ separately for all 14 unique adult tissues and for the 10 non-sex-specific tissues. We also identified the tissue where expression is highest for every gene that has *S_max_* > 100 and where expression was detected for at least one probe in all 4 replicate arrays in that tissue.

## DATA AVAILABILITY

No new data were generated or analyzed in support of this research.

## ACKNOWLEDGEMENTS

We thank Wen-Juan Ma for feedback and suggestions on previous versions of this manuscript. This work was completed in part with resources provided by the Research Computing Data Core at the University of Houston.

---

**Supplemental Figure S1.**
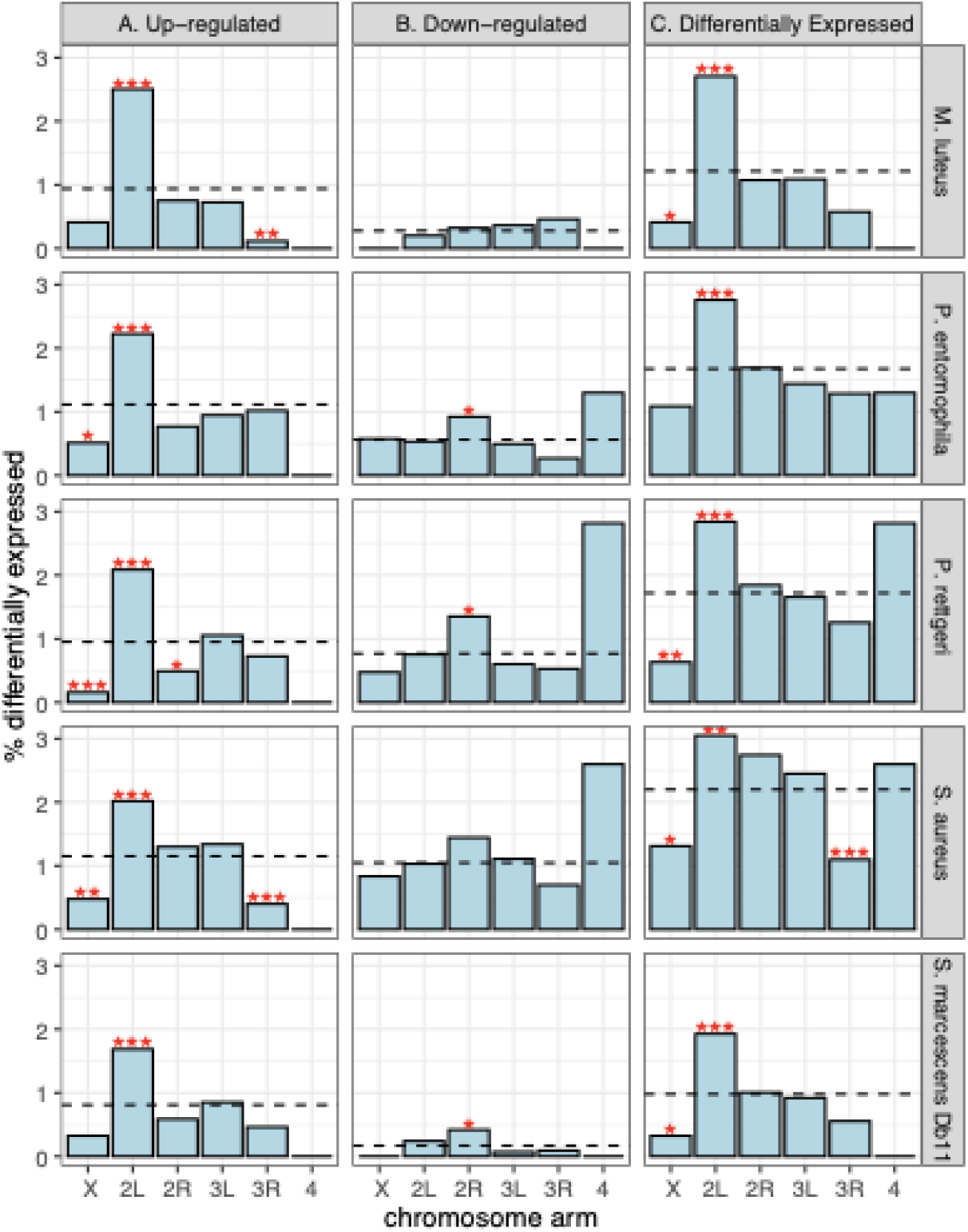
The percent of genes on each chromosome arm that are differentially expressed following infection with one of five bacteria in male *D. melanogaster* is shown. Genes are either (A) up-regulated, (B) downregulated, or (C) differentially expressed (sum of up- and down-regulated). The percent of differentially expressed genes across the entire genome is shown as a dashed line. Asterisks indicate chromosomes where the percent of genes is significantly different from the rest of the genome (\**P*<0.05, \*\**P*<0.005, or \*\*\**P*<0.0005 in Fisher’s exact test).

**Supplemental Figure S2.**
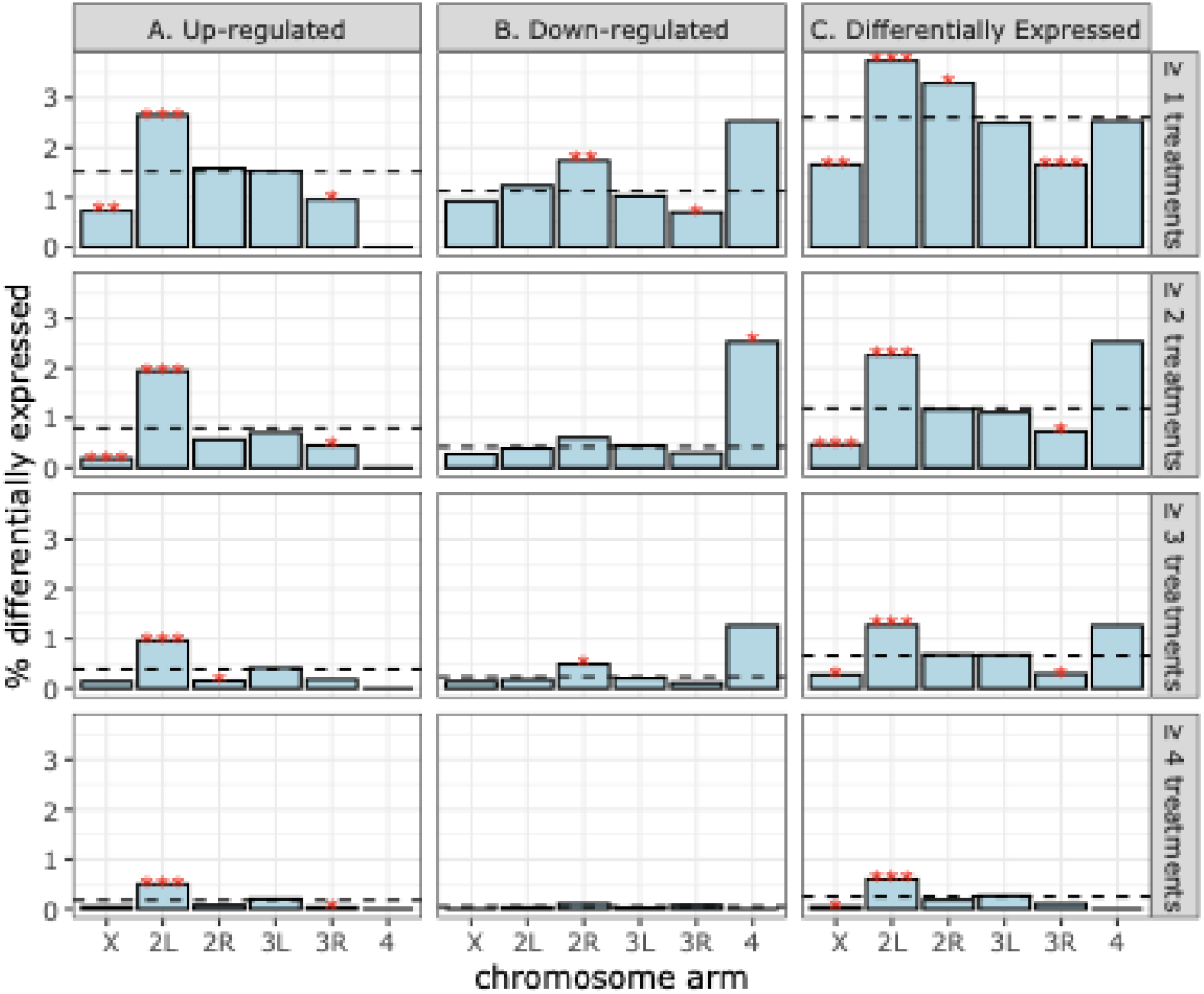
The percent of genes on each chromosome arm that are differentially expressed following infection with at least 1, 2, 3, or 4 different bacteria in male *D. melanogaster* is shown. Genes are either (A) up-regulated, (B) downregulated, or (C) differentially expressed (sum of up- and down-regulated). The percent of differentially expressed genes across the entire genome is shown as a dashed line. Asterisks indicate chromosomes where the percent of genes is significantly different from the rest of the genome (\**P*<0.05, \*\**P*<0.005, or \*\*\**P*<0.0005 in Fisher’s exact test).

**Supplemental Figure S3.**
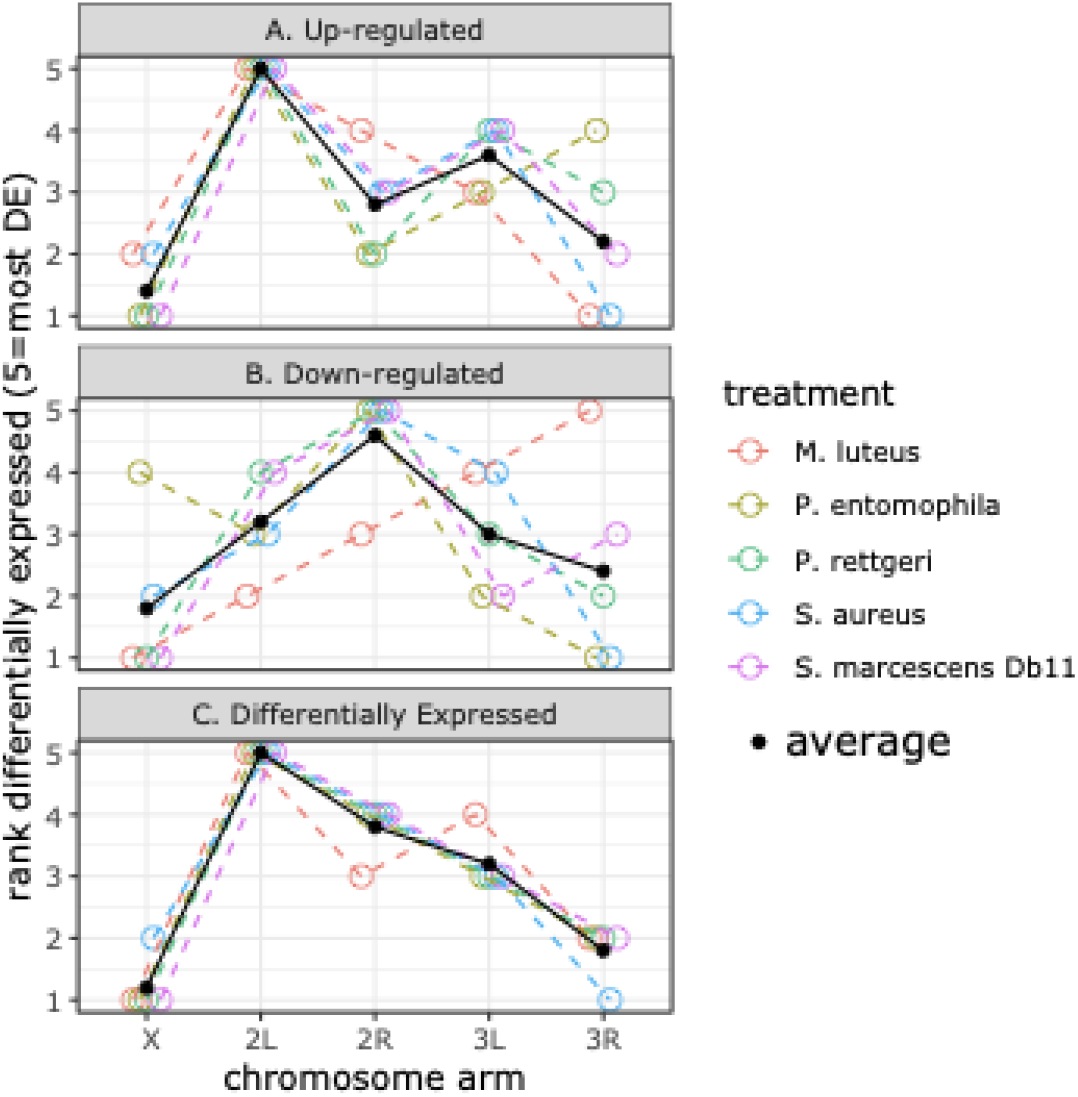
The rank order of chromosome arms is shown according to the percent of differentially expressed genes in each of five different bacterial treatments. The chromosome arm that has the highest percent of differentially expressed genes is ranked as 5, and the chromosome with the lowest percent is ranked as 1. Genes are either (A) up-regulated, (B) downregulated, or (C) differentially expressed (sum of up- and down-regulated). The mean ranks for each chromosome are shown as black dots and solid lines.

**Supplemental Figure S5.**
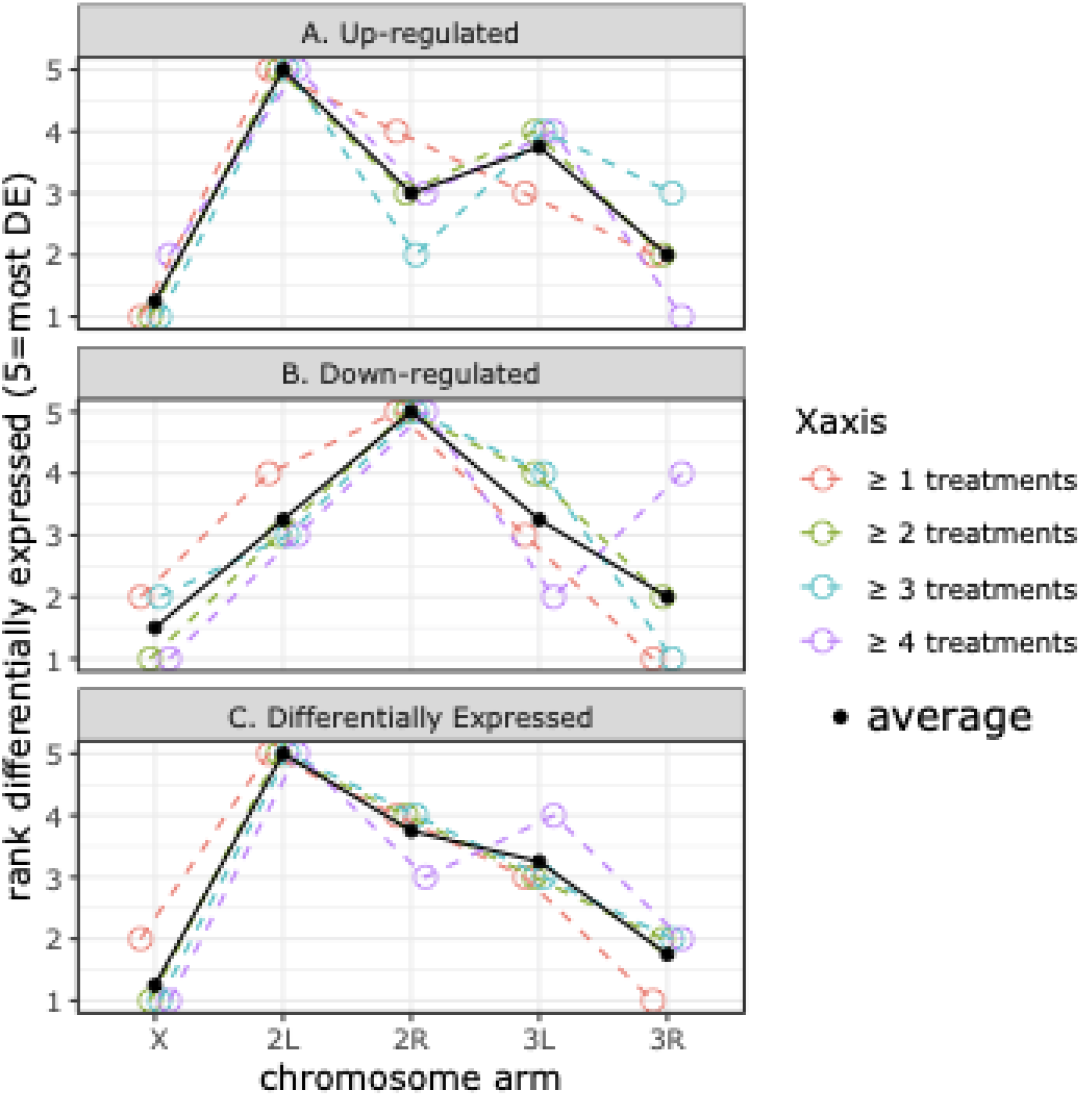
The rank order of chromosome arms is shown according to the percent of differentially expressed genes in at least one, two, three, or four bacterial treatments. The chromosome arm that has the highest percent of differentially expressed genes is ranked as 5, and the chromosome with the lowest percent is ranked as 1. Genes are either (A) up-regulated, (B) downregulated, or (C) differentially expressed (sum of up- and down-regulated). The mean ranks for each chromosome are shown as black dots and solid lines.

**Supplemental Figure S5.**
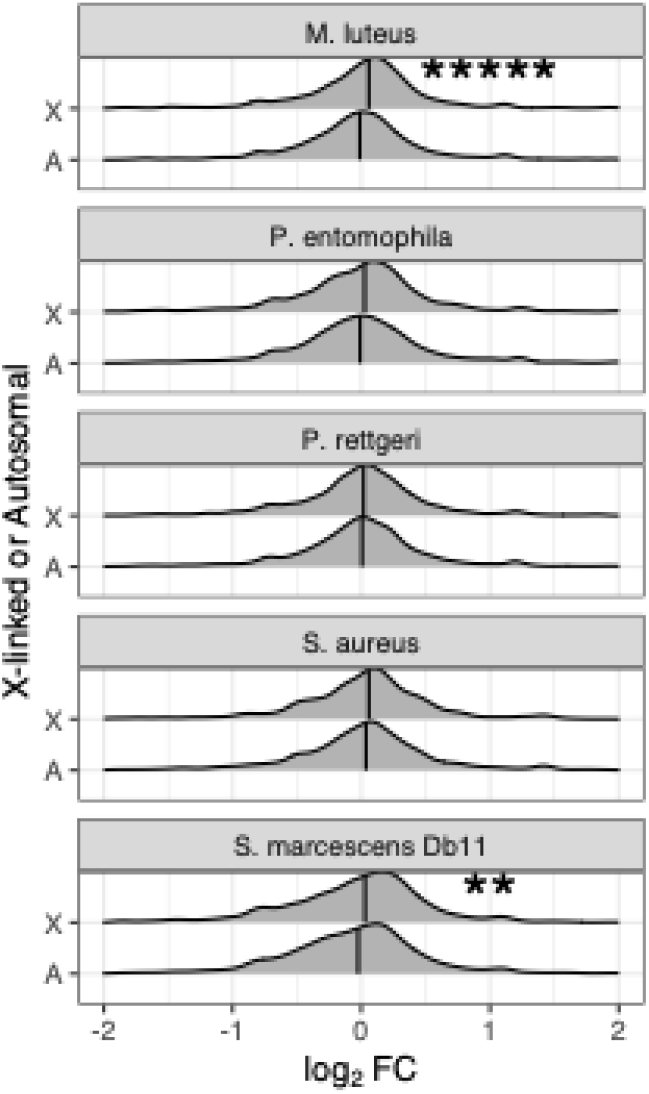
Distributions of log_2_FC following bacterial infection for X-linked (X) and autosome (A) genes in *D. melanogaster* males. Asterisks show significant differences between X and autosomes within a treatment (*\*\**P<0.005; *****P<0.000005; Mann-Whitney test).

**Supplemental Figure S6.**
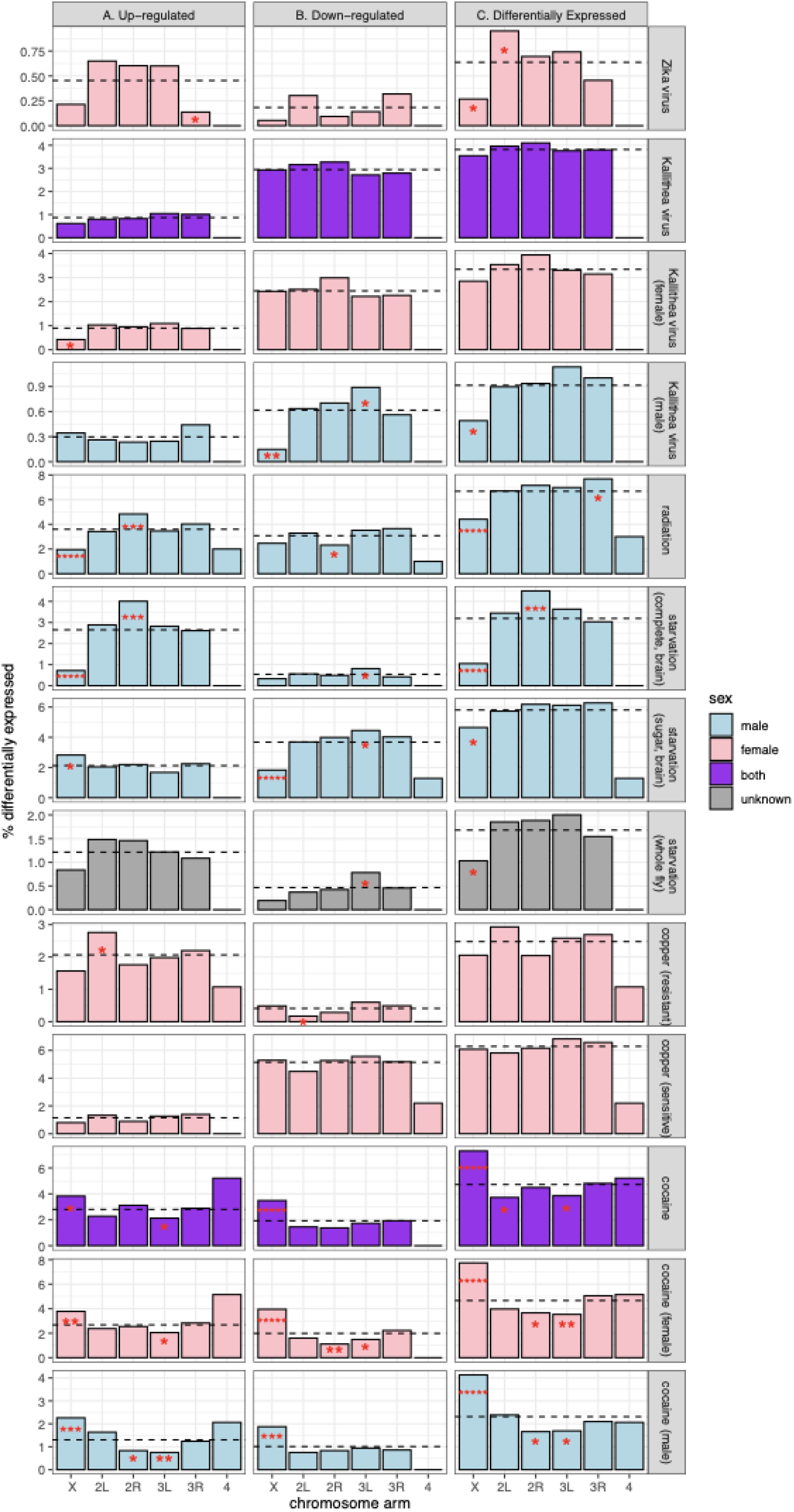
The percent of genes on each chromosome arm that are differentially expressed following viral and abiotic treatments is shown. Bars are colored by the sex of the flies used in each treatment (see legend). Genes are either (A) up-regulated, (B) downregulated, or (C) differentially expressed (sum of up- and down-regulated). The percent of differentially expressed genes across the entire genome is shown as a dashed line. Asterisks indicate chromosomes where the percent of genes is significantly different from the rest of the genome (\**P*<0.05, \*\**P*<0.005, or \*\*\**P*<0.0005 in Fisher’s exact test).

**Supplemental Figure S7.**
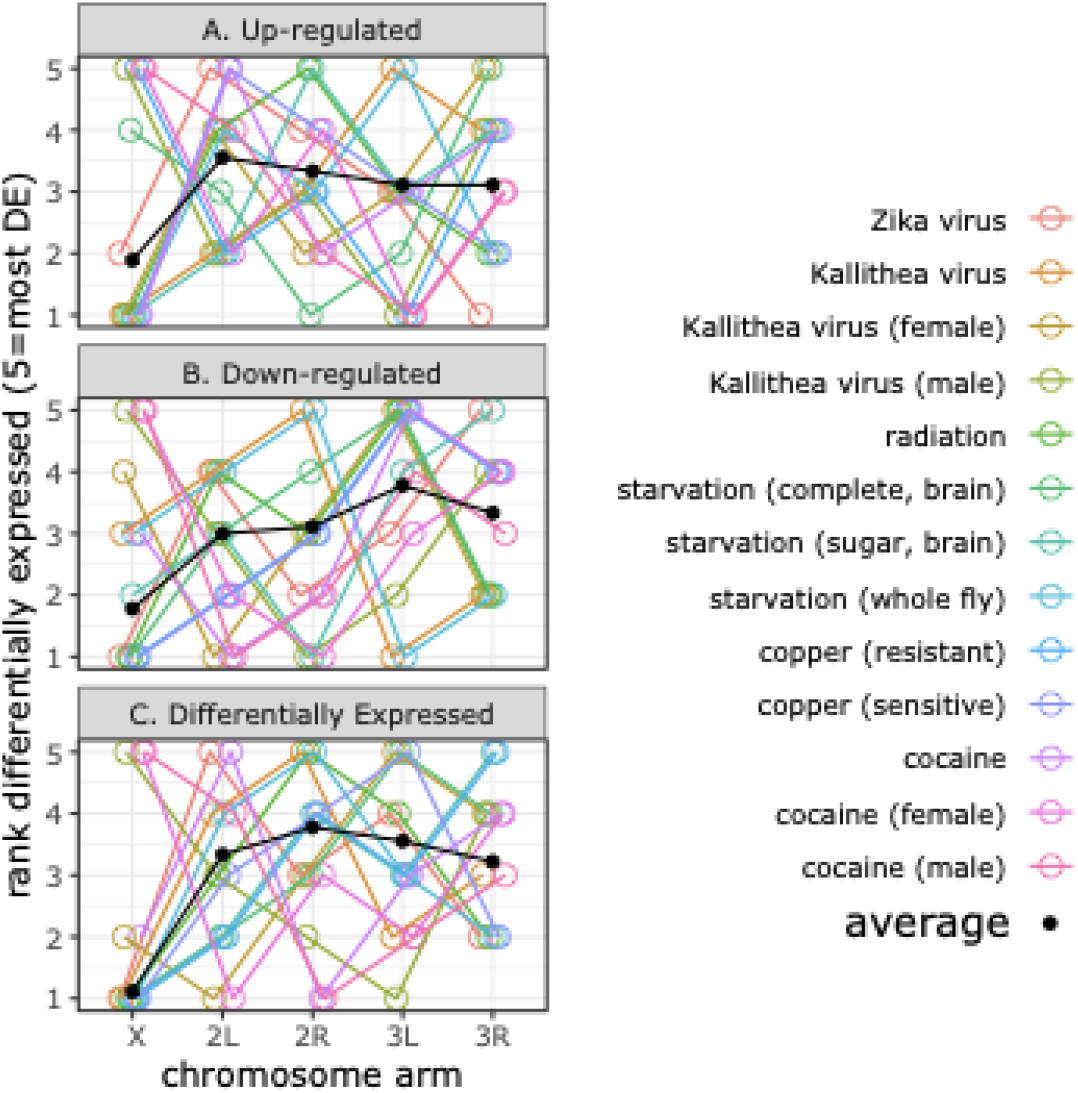
The rank order of chromosome arms is shown according to the percent of differentially expressed genes in viral and abiotic treatments. The chromosome arm that has the highest percent of differentially expressed genes is ranked as 5, and the chromosome with the lowest percent is ranked as 1. Genes are either (A) up-regulated, (B) downregulated, or (C) differentially expressed (sum of up- and down-regulated). The mean ranks for each chromosome are shown as black dots and solid lines.

**Supplemental Figure S8.**
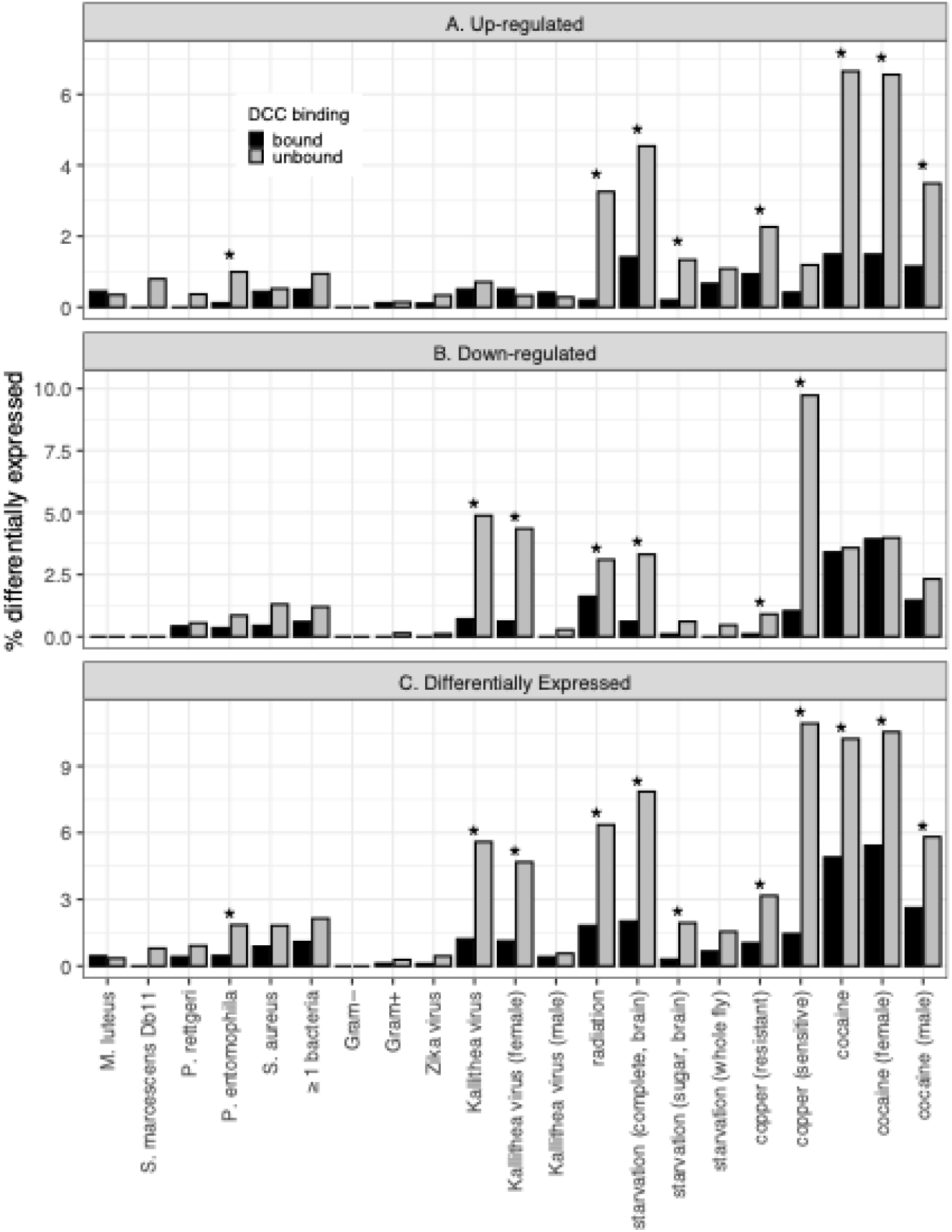
The percent of X-linked DCC-bound (black) and unbound (gray) genes that are up-regulated (A), down-regulated (B), or differentially expressed (C) are shown for each treatment and sample type (in parentheses). Asterisks show a significant difference in the percent between DCC-bound and unbound genes (*P<0.05 in Fisher’s exact test).

**Supplemental Figure S9.**
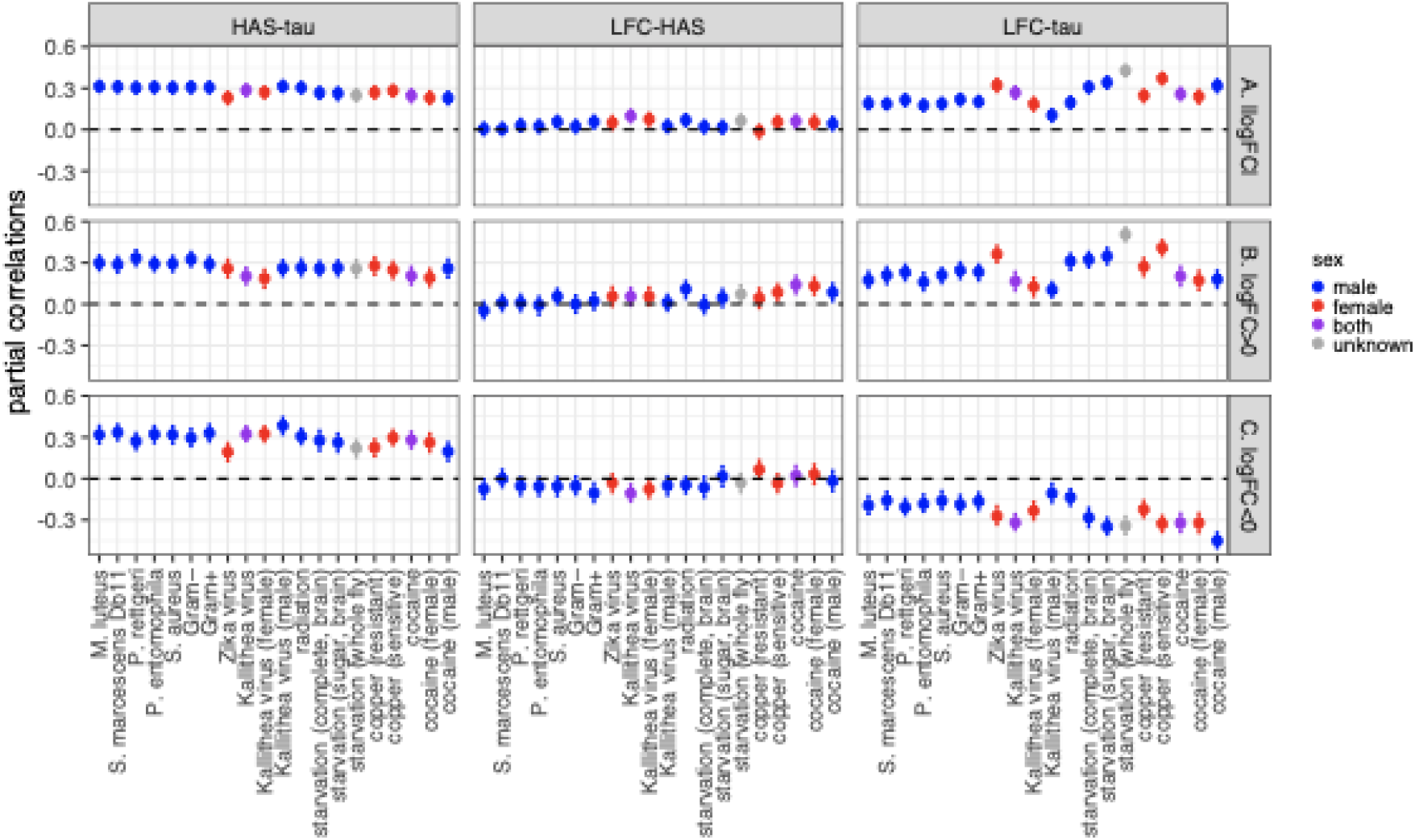
Partial correlations between distance from a DCC high affinity site (HAS), tissue-specificity (tau), and log_2_FC (LFC) are shown for each treatment. Error bars show the 95% confidence interval from 1,000 bootstrap replicate samples of X-linked genes. The analysis was performed on absolute values of log_2_FC of all genes (A), genes with positive log_2_FC>0 (B), or genes with negative log_2_FC<0 (C). Tissue-specificity values (τ) were calculated using all 14 adult tissues. HAS were taken from Alekseyenko et al. (2008).

**Supplemental Figure S10.**
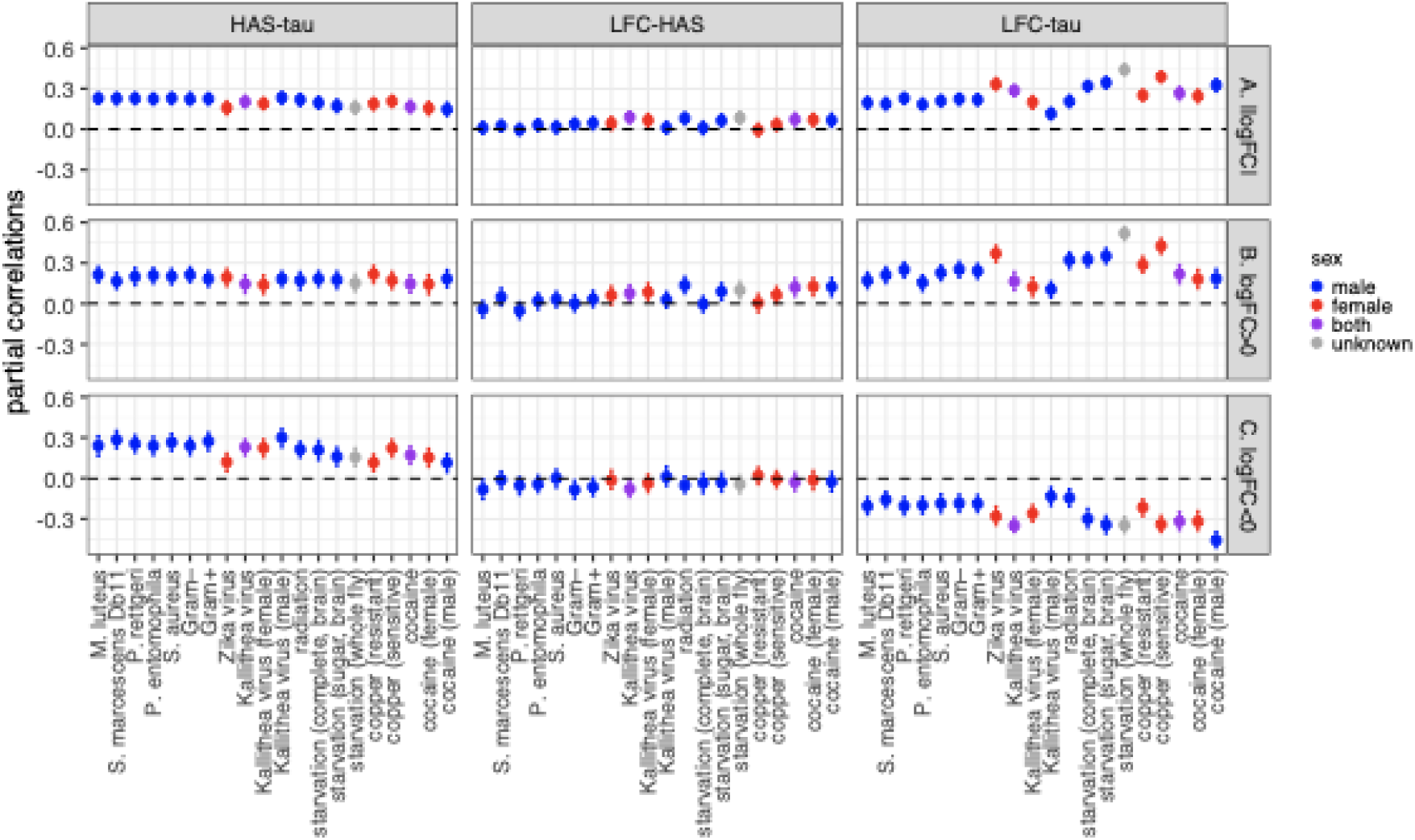
Partial correlations between distance from a DCC high affinity site (HAS), tissue-specificity (tau), and log_2_FC (LFC) are shown for each treatment. Error bars show the 95% confidence interval from 1,000 bootstrap replicate samples of X-linked genes. The analysis was performed on absolute values of log_2_FC of all genes (A), genes with positive log_2_FC>0 (B), or genes with negative log_2_FC<0 (C). Tissue-specificity values (τ) were calculated using all 14 adult tissues. HAS were taken from Straub et al. (2008).

**Supplemental Figure S11.**
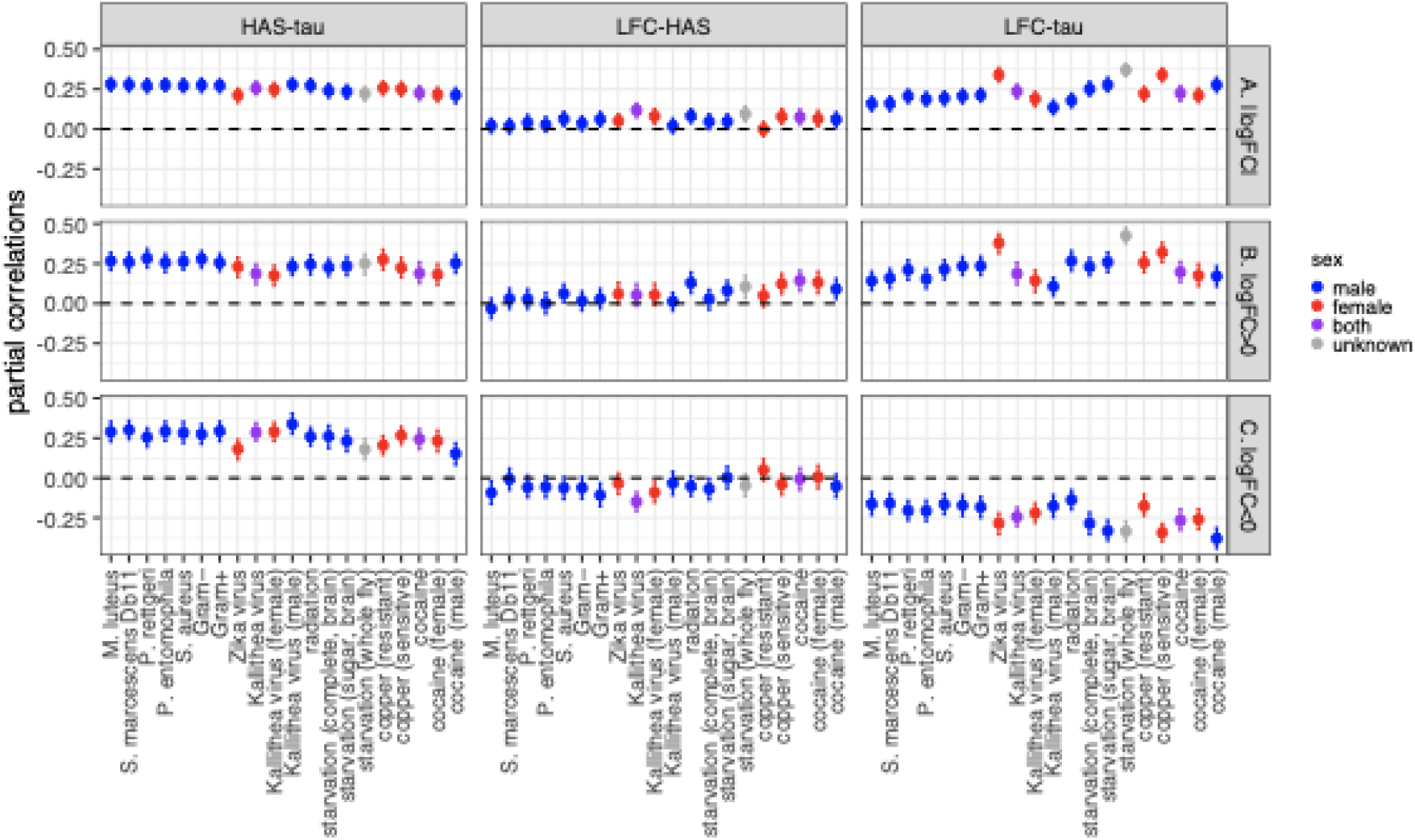
Partial correlations between distance from a DCC high affinity site (HAS), tissue-specificity (tau), and log_2_FC (LFC) are shown for each treatment. Error bars show the 95% confidence interval from 1,000 bootstrap replicate samples of X-linked genes. The analysis was performed on absolute values of log_2_FC of all genes (A), genes with positive log_2_FC>0 (B), or genes with negative log_2_FC<0 (C). Tissue-specificity values (τ) were calculated using 10 non-sex-specific tissues. HAS were taken from Alekseyenko et al. (2008).

**Supplemental Figure S12.**
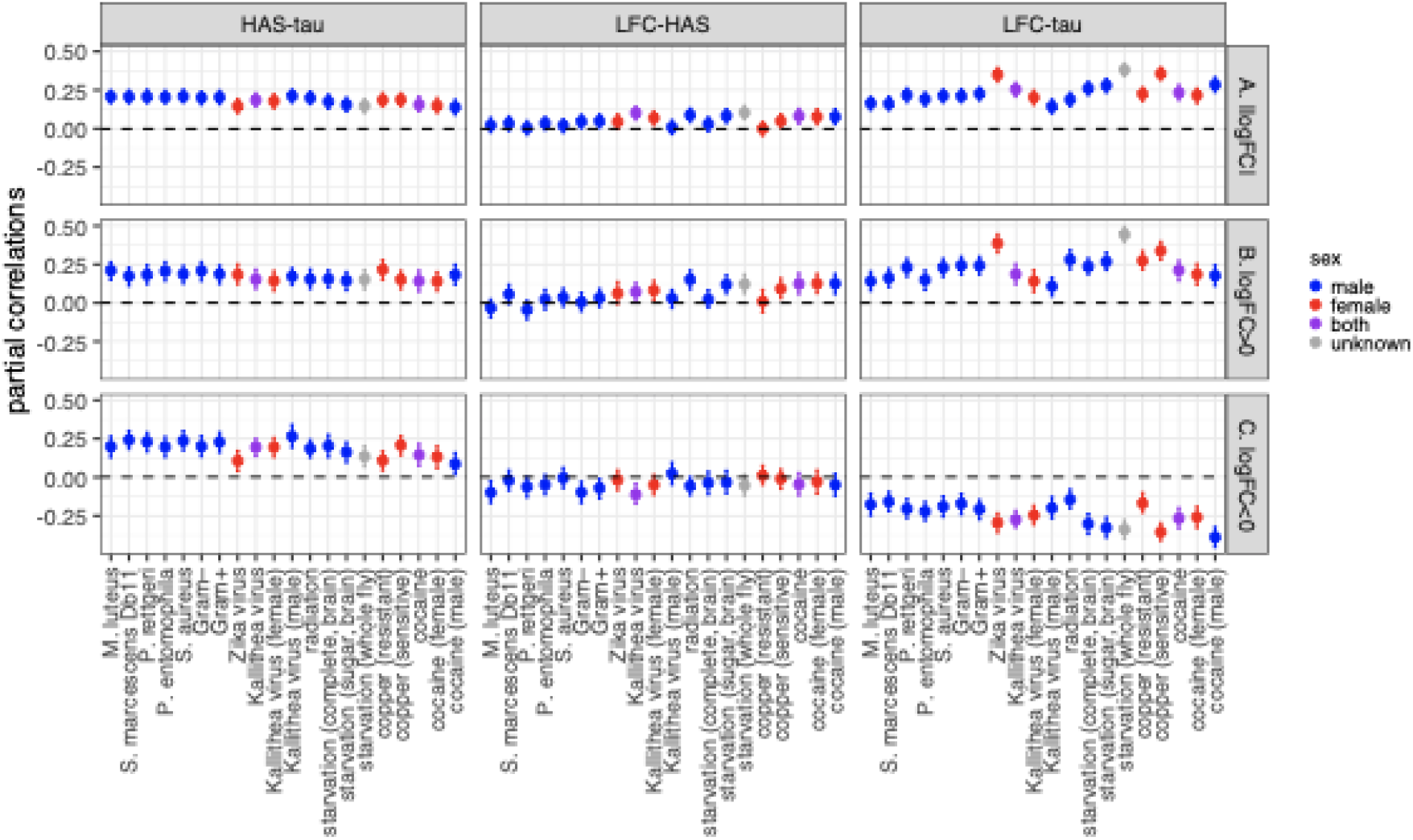
Partial correlations between distance from a DCC high affinity site (HAS), tissue-specificity (tau), and log_2_FC (LFC) are shown for each treatment. Error bars show the 95% confidence interval from 1,000 bootstrap replicate samples of X-linked genes. The analysis was performed on absolute values of log_2_FC of all genes (A), genes with positive log_2_FC>0 (B), or genes with negative log_2_FC<0 (C). Tissue-specificity values (τ) were calculated using 10 non-sex-specific-tissues. HAS were taken from Straub et al. (2008).

**Supplemental Figure S13.**
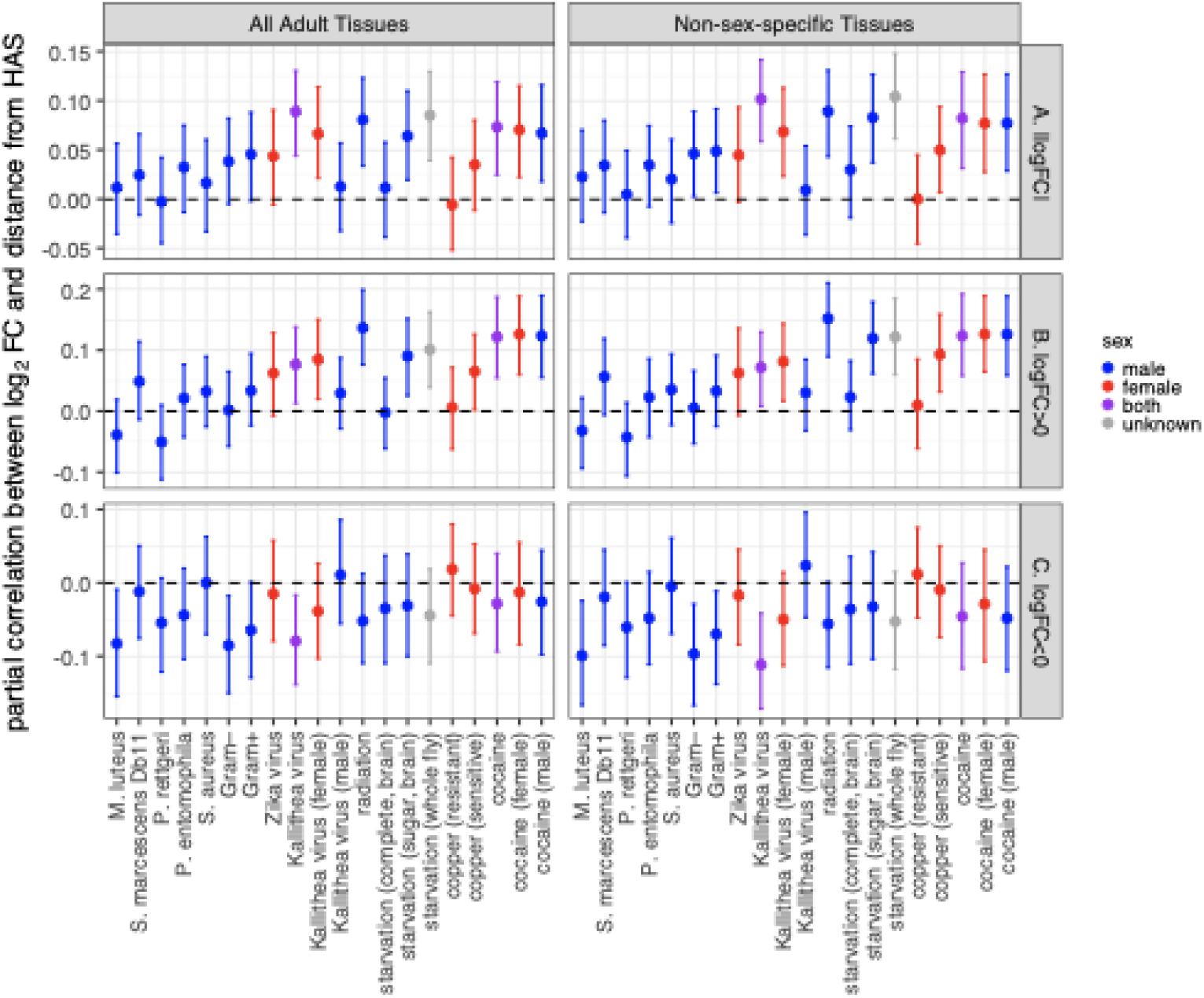
Partial correlations between log_2_ fold-change in expression between treatment and control (log_2_FC) and distance from a dosage compensation complex high affinity site (HAS). Partial correlations were calculated based on rank order correlations between log_2_FC, distance from an HAS, and tissue expression breadth. Each dot shows the partial correlation between log_2_FC and distance from an HAS, with the error bars representing 95% confidence intervals from 1,000 bootstrap replicates of the data. The X-axis shows the specific treatment. Dots and error bars are colored based on the sex of the flies used in the experiment (see legend). HAS were obtained from the Straub *et al*. (2008) data set. Expression breadth was calculated using microarray data from either 14 unique adult tissues (left) or 10 adult tissues that are not sex-specific (right). Partial correlations are plotted with |log_2_FC| values for all genes (A), only genes with log_2_FC>0 (B), and only genes with log_2_FC<0 (C).

